# The HIV-1 provirus excised by a single CRISPR/Cas9 RNA guide persists in the host cell and may be reactivated

**DOI:** 10.1101/2020.11.16.384180

**Authors:** Michele Lai, Eyal Maori, Paola Quaranta, Giulia Matteoli, Fabrizio Maggi, Marco Sgarbanti, Stefania Crucitta, Simone Pacini, Ombretta Turriziani, Giulia Freer, Guido Antonelli, Jonathan L. Heeney, Mauro Pistello

**Affiliations:** Retrovirus Center and Virology Section, Department of Translational Research and New Technologies in Medicine and Surgery, University of Pisa, Pisa, Italy; Laboratory of Viral Zoonotics, Department of Veterinary Medicine, University of Cambridge, Cambridge, United Kingdom; Virology Unit, Pisa University Hospital, Pisa, Italy; National Institute of Health, Rome, Italy; Pharmacology Unit, Department of Clinical and Experimental Medicine, University of Pisa, Italy; Hematology Unit, Department of Clinical and Experimental Medicine, University of Pisa, Italy; Laboratory of Virology and Pasteur Institute-Cenci Bolognetti Foundation, Department of Molecular Medicine, Sapienza University of Rome, Rome, Italy

**Keywords:** CRISPR/Cas9, Gene therapy, Endonucleases, Gene editing, HIV, Latent reservoir, Integrase, Tat, Rev, J-Lat

## Abstract

Gene editing may be used to cut out the human immunodeficiency virus type-1 (HIV-1) provirus from the host cell genome and eradicate infection. Here, using cells acutely or latently infected by HIV and treated with long terminal repeat-targeting CRISPR/Cas9, we show that the excised HIV provirus persists for a few weeks and, by means of HIV Integrase, rearranges in circular molecules. Circularization and integration restore proviral transcriptional activity that is enhanced in the presence of exogenous Tat and Rev or tumor necrosis factor-α, respectively, in acutely or latently infected cells. Although confirming that gene editing is a powerful tool to eradicate HIV infection, this work highlights that, to achieve this goal, the provirus has to be cleaved in several pieces and the infected cells treated with antiviral therapy before and after editing.

## Introduction

The highly active anti-retroviral therapy (HAART) efficiently abates human immunodeficiency virus type-1 (HIV-1) replication and has transformed a deadly infection into a chronic illness. Unfortunately, HAART does not provide a cure. By stalling viral replication, HAART halts HIV spread to other cells but allows HIV to persist and reactivate at any time. Clustered regularly interspaced short palindromic repeats/Cas9 (CRISPR/Cas9), a technique that is changing paradigms and expectancies to cure genetic maladies^1^, holds promises to provide a cure for HIV too^2^. CRISPR/Cas9 cuts out the integrated HIV-1 genome (provirus) from the host cell genome^3^ and has proved effective to eliminate, within certain limits, infection *in vitro* and *in vivo*^4^.

The HIV long terminal repeats (LTRs) are highly conserved among HIV strains and comprise sequence domains recognized by cellular and viral proteins driving viral replication. Initial studies to eliminate HIV infection were performed using a single CRISPR/Cas9 RNA guide recognizing a sequence domain present in both LTRs of nearly all strains^3^. This approach, while effective at curing some infected cells, has been recently demonstrated to facilitate virus escape^5, 6, 7^ through the ensuing non-homologous end joining (NHEJ) repair mechanism^8^. It has also been shown that CRISPR/Cas9 alone is not always sufficient to eliminate HIV infection^4^. At the moment, however, CRISPR/Cas9 is the most effective gene editing method to cure HIV-infected cells, it has proved safe for human cells and animal models and has, therefore, good prerequisites for clinical use^9,10, 11^. The rationale of the present study is based on previous observations showing that most linear or circularized reverse transcribed HIV RNA genome (cDNA) produced during viral replication does not integrate^12,13^. Such unintegrated viral molecules are thought to either be destroyed, or aid productive infection through expression of several genes^14^, or have a second chance of integrating through complementation or rearrangement events^15^. The aim of this study is to investigate the fate of the provirus once excised from the host cell genome. To this purpose, and taking into account current difficulties to deliver multiple RNA guides per cell *in vivo*^16^, we used a single CRISPR/Cas9 RNA guide targeting both LTRs. Experiments were conducted in human embryonic kidney 293 (293T) cells bearing integrated, labeled HIV molecular clones and then extended to human T-lymphoid cells actively replicating HIV-1, as occurs during acute infection, or latently infected J-lat cells, as observed in the asymptomatic phase. The results show that the excised provirus persists in the nucleus for a prolonged period of time; depending on the number of copies per cell, it closes as single or rearranged inter-molecular circular elements that can form complete LTRs again with the aid of HIV Integrase (IN), thereby reducing the efficiency of HIV eradication by gene editing. We also show that pretreatment with Raltegravir (RAL) and Efavirenz (EFV) prevents such events. Circular forms generated by inter-molecular joining exhibit functional LTRs that drive viral transcriptional activity and respond to exogenous Tat and Rev, as occurs naturally during superinfection.

## Results

### CRISPR/Cas9 treatment efficiently excises the HIV-1 provirus and triggers non-homologous end-joining mechanisms to repair cellular DNA breaks

Human 293T cells were transduced with NL4-3/Luc or NL4-3/GFP, two HIV-1 NL4-3 reporter lentiviral particles obtained by pseudotyping with glycoprotein of the vesicular stomatitis virus (VSV-G). NL4-3/Luc is derived from pNL4-3 Luc.R-E- and encodes luciferase (Luc)^17^. NL4-3/GFP is derived from pNL4-3 ΔEnv EGFP and expresses the green fluorescent protein (GFP) as Env-GFP fusion protein that is retained in the endoplasmic reticulum^17^. The NL4-3-based constructs express their genes and GFP or Luc reporter genes under the control of 5’-LTR, but they are defective of *env* and, therefore, do not produce infectious particles unless *env* is provided *in trans*. The transduced cells were maintained in culture for two weeks to obtain stably integrated lines. Cells were then transfected with a plasmid expressing CRISPR/Cas9 and Puromycin resistance gene, and HIV-1 specific (T5) or scramble (SC) guide RNAs (gRNAs). The T5 gRNA targeted a highly conserved region between TATA box and NF-κB binding site in LTRs (Supplementary Information). CRISPR/Cas9-transfected cells were selected using high dose Puromycin, which eliminated non-transfectants within three days of treatment (data not shown) and were monitored for reduction of Luc activity. As shown in Fig. 1a, in cells treated with CRISPR/Cas9 and T5 gRNA, Luc activity dropped abruptly to less than 25% between day 2 and 3 after transfection and declined to nearly undetectable levels thereafter. To rule out that Luc reduction was caused by cell death or arrest of cell growth as a consequence of transfection, we performed a cell viability assay (WST-8) 48 h post-transfection. As shown in Fig. 1b, no differences were observed amongst cell populations thus demonstrating that transfection did not damage cells and Luc reduction was indeed due to HIV editing.

**Fig. 1.**
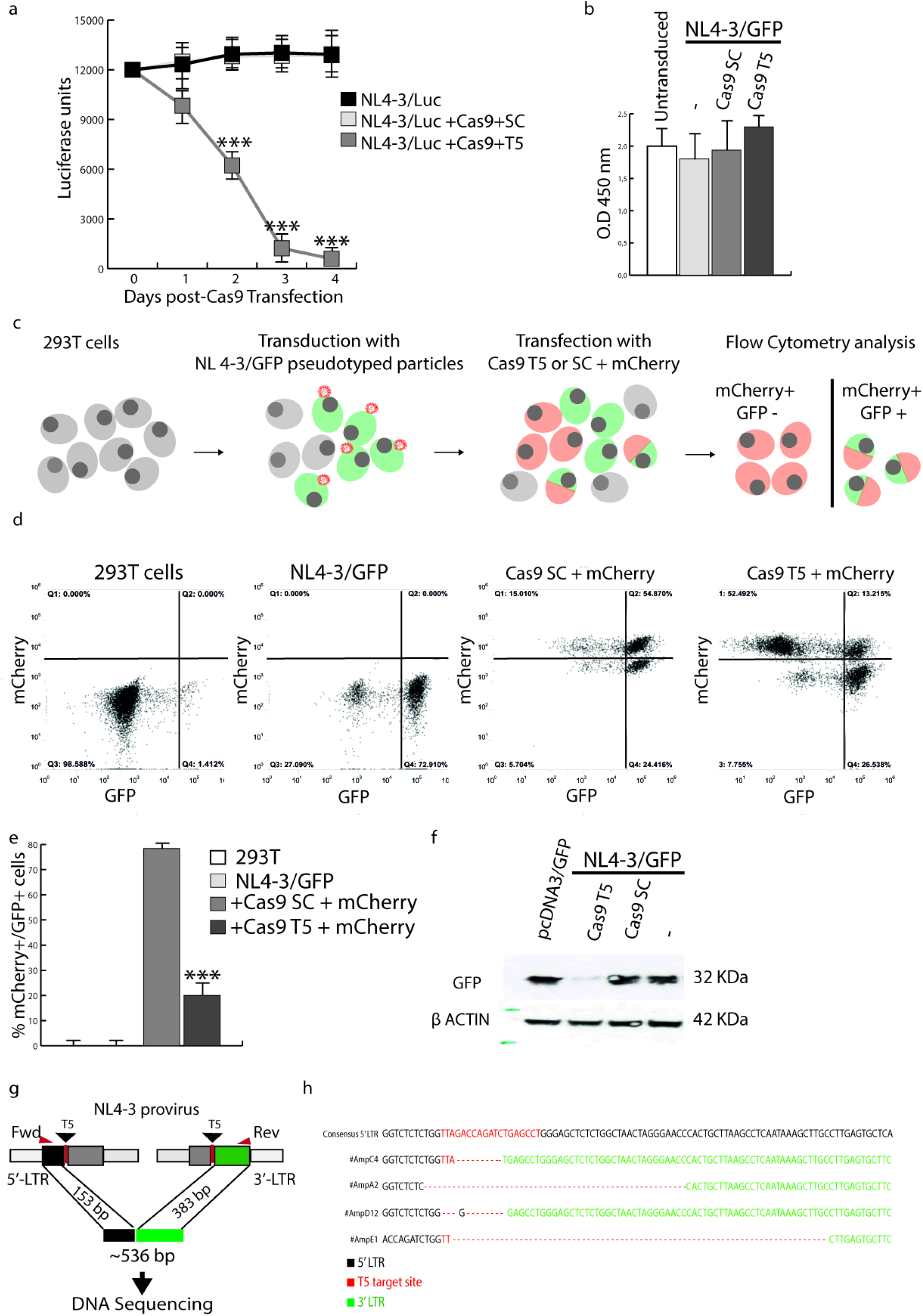
CRISPR/Cas9 efficiently cleaves the HIV-1 provirus in 293T cells transduced with NL4-3/GFP or NL4-3/Luc. **a,** 293T cells that had been transduced with NL4-3/Luc were transfected with CRISPR/Cas9 and either scrambled (SC) or HIV-1 specific (T5) gRNAs and analyzed for Luc expression at day 0, 1, 2, 3, 4. Decrease in Luc expression in T5 gRNA-treated cells compared to untreated or SC gRNA-treated cells reached statistical significance at day 2 (*p*<0.001). Standard deviation (SD) was calculated from three independent experiments. **b,** Cell viability was assessed by WST-8 48 hours after CRISPR/Cas9 transfection and showed no significant difference between cells transfected with NL4-3/Luc alone or in combination with Cas9+SC or +T5 gRNAs, and untreated cells. Shown are the means of three independent experiments with 6 technical replicates (n = 3; m = 6) and SD. Statistical analyses were performed using the Student’s T test. **c,** Schematic of the flow cytometry analysis of GFP expression by 293T cells either not transduced or that had been previously transduced with NL4-3/GFP pseudotyped with VSV-G, and left as such (NL4-3/GFP) or then transfected with the SC-or T5 gRNA-containing CRISPR/Cas9 mCherry plasmid. **d,** Flow cytometry analysis of cells described in **c** was performed at day 4 post-transfection, showing that T5 significantly decreased, but did not ablate, GFP expression by the provirus. **e,** Statistical analysis of the values obtained in **d**. Shown are the means of three independent experiments with biological triplicates (n = 3; m = 3) and the SD. Statistical analyses were performed using the Student’s T test (*** = *p*<0.001). **f,** Western blot analysis of GFP protein levels at day 7 in lysates from 293T cells first transduced with NL4-3/GFP then either left as such (−) or transfected with Cas9+SC or +T5 gRNAs. A control of cells transfected with a GFP coding plasmid (pcDNA3/GFP) is included. **g,** Localization of primers used to amplify the DNA fragments encompassing the CRISPR/Cas9 T5 cleavage sites. **h,** Sequence data of four amplicons sequenced at the LTR junctions and blasted against pNL4-3.

To better understand how CRISPR/Cas9 editing affected HIV transcription, we transduced 293T cells with NL4-3/GFP and, 2 days later, we either left them as such (NL4-3/GFP) or transfected them with SC-or T5 gRNA-containing CRISPR/Cas9 and red fluorescent protein (mCherry) plasmid, to follow edited cells (Fig.1c). Consistently with the Luc data, CRISPR/Cas9 treatment caused sharp reduction in GFP expression, so that the number of mCherry+/GFP+ cells dropped to about 10% four days after transfection (Fig. 1d and 1e). In addition, GFP was nearly undetectable by Western blot analysis at day 7 (Fig. 1f). No changes were observed after treatment with CRISPR/Cas9 combined with SC gRNA (Fig. 1a, b, d-f). The criteria used for target selection, T5 and SC gRNA sequences, and CRISPR reaction conditions are provided in Supplementary Information.

To investigate if excision events had occurred and how cellular DNA repair had resolved the editing, we obtained individual clones of CRISPR/Cas9-transfected NL4-3/Luc cells, by limiting dilution five days post-CRISPR/Cas9 treatment. Amplification was performed using a semi-nested PCR with primers annealing upstream and downstream the T5 target site (Fig. 1g) and expecting an amplicon of roughly 536 base pairs (bp) as a result of mere joining of edited genomic DNA ends with no rearrangements in-between (Fig. 1g). As expected, PCR yielded amplicons of about 500 bp that were cloned and sequenced at random. As shown in Fig. 1h, most junctions were achieved by NHEJ and presented with deletions of DNA fragments of variable lengths.

### The excised HIV-1 provirus persists and circularizes in NL4-3/Luc-transduced CRISPR/Cas9-transfected 293T cells

To investigate the fate of HIV-1 provirus after excision and determine whether the proviral fragments circularize, genomic DNA was extracted from NL4-3/Luc-transduced 293T cells at various days post-CRISPR/Cas9 and T5 or SC gRNA transfection and digested with an ATP-dependent DNA exonuclease to eliminate linear DNA fragments. The digested DNA was first checked for residual linear DNA by *β-globin* amplification (Fig. 2a) and then amplified for *gag* and *pol* (primer sequences available in Supplementary Information). As determined by agarose gel electrophoresis, HIV sequences were found in the exonuclease-digested DNA samples up to 14 days post-CRISPR/Cas9 T5 gRNA treatment (Fig. 2b). In contrast, no *gag* or *pol* sequences were found in the DNA samples treated with SC gRNA and digested with the exonuclease (data not shown). Since PCR concatemer analysis was performed in a population of dividing cells and the concatemers were likely to be less and less during subsequent cell mitosis, it is not known whether the PCR signal disappeared because DNA molecules were eventually degraded or progressively diluted out. Anyhow, these results suggest that, upon excision, the provirus persists for at least a couple of weeks, most likely in circular or similar DNA exonuclease-resistant forms.

**Fig. 2.**
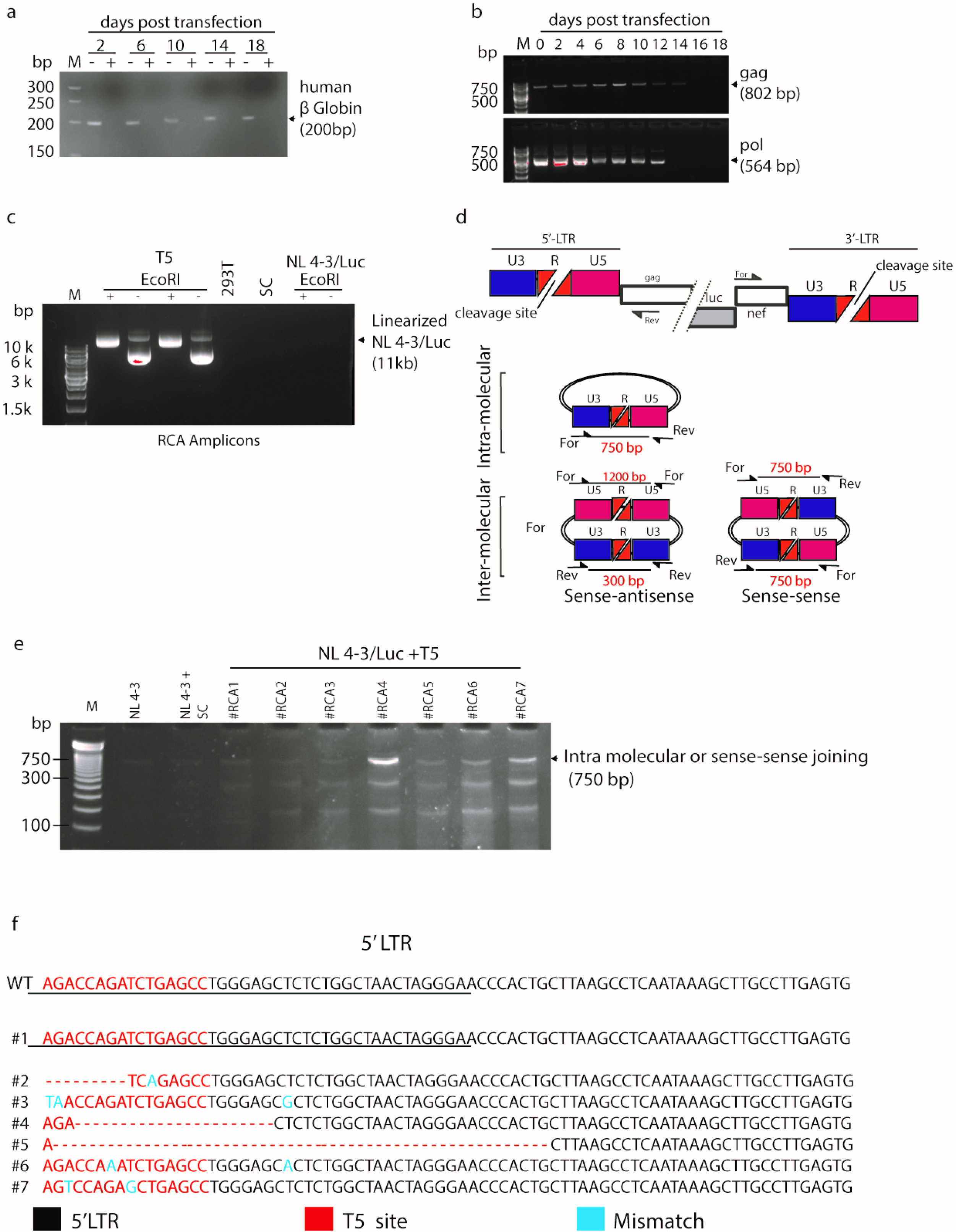
The excised HIV-1 provirus persists and circularizes after CRISPR/Cas9-transfection. **a,** PCR analysis of *β-globin*gene in genomic DNA samples from NL4-3/Luc-transduced 293T cells extracted at day 2, 6, 10, 14 and 18 post CRISPR/Cas9 transfection, before (−) and after (+) ATP-dependent DNA exonuclease digestion. M: Molecular weight marker. **b,** PCR analysis of *gag* and *pol* sequences of genomic DNA treated as in **a** with ATP-dependent DNA exonuclease digestion. **c,** Rolling circle amplification (RCA) of DNA samples extracted from NL4-3/Luc-transduced at day 10 post-CRISPR/Cas9 and T5 or SC gRNAs transfection. This technique allows selective amplification of circularized DNA as concatemers, requiring digestion with a single cutter to obtain full length fragments. RCA amplicons were electrophoresed as such (−) or after *EcoR*I digestion (+), which cuts NL4-3/Luc once in *pol*. 293 T were used as a negative control. **d,** Schematic of circular molecules of HIV-1 provirus formed by a single molecule or two molecules bound together in sense-to-sense or sense-to-antisense orientation and size of the amplicons of interest written in red and generated by the primers indicated. **e,** PCR fragments obtained from RCA amplicons from 7 different experiments (#RCA1-7) using the primers shown in Fig. 2d. **f,** Alignment of LTR sequencing of the 750-bp fragments obtained from #RCA1-7 and retrieved from the agarose gel of Fig. 2e. The red sequences indicate the T5 gRNA annealing site. Dashes indicate base deletions and blue letters denote nucleotide mismatches compared to wild-type pNL4-3 sequence.

To explore such a conclusion, we repeated the above experiment with NL4-3/Luc-transduced 293T cells that were transfected with CRISPR/Cas9+T5 and then examined for the presence of circular DNA molecules. To this aim, five replicas of such cells were transduced with NL4-3/Luc, propagated for two weeks and treated with CRISPR/Cas9 and T5 gRNA (three replicas), SC gRNA (one replica), or left untreated (one replica). Cells were propagated for 10 days in the presence of Puromycin to enrich in CRISPR-transfected cells and processed to extract whole DNA. This DNA was exonuclease-digested, checked for *β-globin* amplification (data not shown) and subjected to rolling circle amplification (RCA) to enrich in circular DNA molecules. This technique allows selective amplification of circularized DNA as concatemers, requiring digestion with a single cutter to obtain full length fragments. Fig. 2c shows the RCA amplicons electrophoresed as such or after digestion with *EcoR*I, which cuts pNL4-3/Luc once. Upon transfection with T5 gRNA, one sample produced a smear and was not analyzed any further, while two samples contained RCA amplicons of large size that, upon *EcoR*I digestion, yielded a discrete band (Fig. 2c). Of note, this predominant band had a size compatible with full-length HIV-Luc genome. Conversely, normal 293T cells or cells transduced with NL4-3/Luc and transfected or not with CRISPR/Cas9 + SC gRNA had negligible amounts of circular DNA molecules (Fig. 2c). In all, this experiment indicates that the amount of circular HIV molecules is increased in T5 gRNA-transfected cells, possibly originating from the excised provirus. This experiment also demonstrates that the circularized provirus persisted for a prolonged period of time and, as judged by the size of *EcoR*I fragments, did not undergo large deletions.

To evaluate whether circularization occurred intra-or inter-molecularly, *EcoR*I-digested RCA products of approximately 11 Kb were extracted from agarose gel, divided into four replicas per fragment, diluted to 1 ng and amplified in their LTR junctions. To this aim, we designed forward (Fwd) and reverse (Rev) primers annealing to *luc/nef* and *gag* regions (Fig. 2d), respectively, and generating amplicons the size of which depended on whether circularization had occurred within a single excised provirus (intra-molecular circularization) or between two or more molecules (inter-molecular circularization). In particular, as shown in Fig. 2d, a single, circularized molecule or two HIV proviral molecules bound in sense-sense orientation would yield an amplicon of 750 bp. Conversely, two molecules bound in sense-antisense orientation would yield two amplicons. The first, obtained by extending two Fwd primers, would be 300 bp, the second, obtained with two Rev primers, would be 1200 bp (Fig. 2d). Because circular DNA generated by sense-sense inter-molecular joining has the potential to recreate full-length LTRs (Fig. 2d), we used PCR conditions privileging amplification of shorter fragments. Cells transfected with SC gRNA produced a faint band of 750 bp, as expected for a circular monomer; cells treated with T5 gRNA yielded more amplicons, two of which compatible with the estimated 750 and 300 bp (Fig. 2e). The 750-bp amplicons shown in Fig. 2e were then retrieved from agarose gel and sequenced. Most of them had In/Dels. Of note, one of them had a wild-type sequence demonstrating that intra-molecular or inter-molecular sense-sense orientation could recreate wild-type LTRs (Fig. 2f).

### The excised HIV-1 provirus can generate concatemers through inter-molecular joining

To further confirm that circularization of excised proviruses can also occur through binding of two or more molecules in sense-sense orientation, we devised the approach shown in Fig. 3a. Using the NL4-3/Luc backbone, to prepare VSV-G pseudotyped viral vectors, we constructed two different HIV-1 molecular clones named NL4-3/Luc/Ori and NL4-3/Luc/KanR (Fig. 3b). NL4-3/Luc/Ori contained the low copy bacterial origin of replication SC101, and NL4-3/Luc/KanR encoded the Kanamycin resistance gene. The genes were cloned in the same restriction site within Δ*env* (Fig. 3b). This would allow to identify and select inter-molecular concatemers, which may generate *in vitro* after CRISPR/Cas9 excision, because inter-molecular concatemers between the two constructs (Ori+KanR) could be used as plasmids to transform bacteria that would only grow with Ori *and* KanR on Kanamycin-containing agar plates. The respective VSV-G-pseudotyped particles were first tested for competence for transduction. Both NL4-3/Luc/Ori and NL4-3/Luc/KanR particles had slightly reduced transduction capacity compared to NL4-3/Luc, but it was deemed sufficient to perform the subsequent experiments (Fig. 3c). As shown in Fig. 3a, 293T cells were co-transduced with NL4-3/Luc/Ori and NL4-3/Luc/KanR, then cultivated for two weeks to obtain stable transduction. Cells were cloned by limiting dilution and probed to select for double-positive clones. Screening was performed with an upstream primer annealing to pNL4-3 and downstream primers annealing to Ori or KanR. Ori and KanR primers were also designed to yield amplicons of about 220 and 280 bp, respectively, to discriminate both clones by agarose gel electrophoresis (Fig. 3d). Double-positive cell clones were then pooled, treated with CRISPR/Cas9 and T5 gRNA, or SG gRNA, to excise both NL4-3/Luc/Ori and NL4-3/Luc/KanR, and cultivated for two weeks in the presence of Puromycin to select for CRISPR/Cas9 transfectants. Whole cellular DNA was then extracted and treated with DNA exonuclease to eliminate the linear fragments (Fig. 3a). To check if the excised NL4-3/Luc/Ori and NL4-3/Luc/KanR had formed inter-molecular concatemers, which we named NL4-3/Luc/KanR/Ori, we took advantage of *KanR,* the Kanamycin resistance gene, and SC101, the bacterial origin of replication, to select and expand inter-molecular concatemers in bacterial cells. DNA exonuclease-digested cellular DNA was thus used to transform bacterial cells which were grown in the presence of Kan (Fig. 3a). From 200 ng of cellular DNA, measured before exonuclease digestion, we obtained 30-50 bacterial colonies. Two hundred ng of cellular DNA is roughly equivalent to 28,000 cells. Considering that this amount was used to transform 10^8^ bacterial cells with a transformation efficiency below 10% for plasmids sizing 20 Kb, we estimate that an inter-molecular concatemer was present in about 1 out to 1,000 cells.

**Fig. 3.**
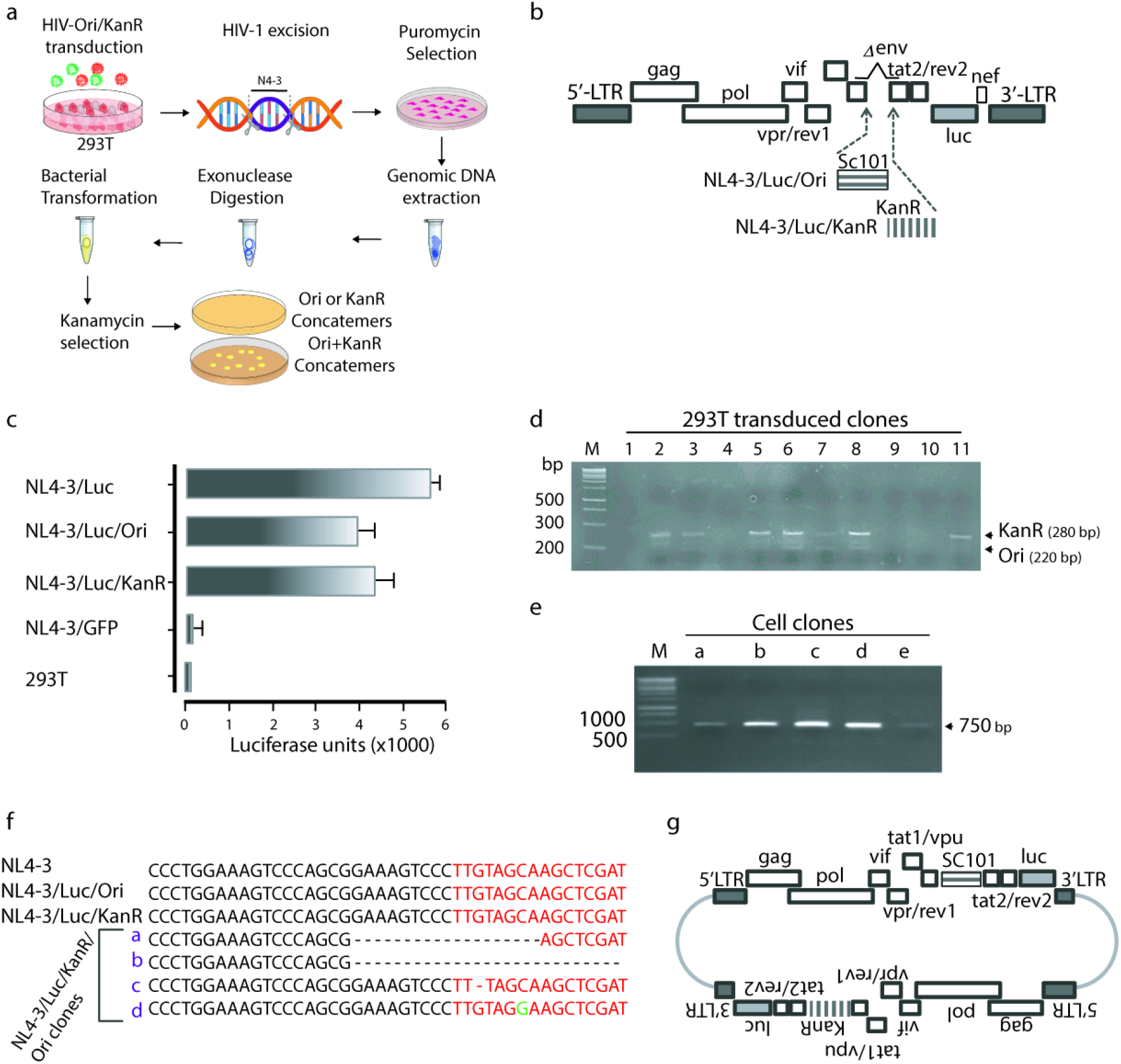
HIV intermolecular concatemers can be isolated in large scale. **a,** Schematic of the approach devised to select and identify inter-molecular concatemers. 293T cells were transduced at 5 MOI with both HIV-Ori and HIV-KanR. A week later, their HIV-1 provirus was excised by CRISPR/Cas9 + T5 gRNA and the edited cells were selected by Puromycin. Then, their genomic DNA was extracted, digested by ATP-dependent exonuclease to eliminate linear DNA and used to transform bacteria. The recombinant bacteria were then selected on Kanamycin-containing agar plates, where only intermolecular joining brought about growing colonies. **b,** Graphic map of NL4-3/Luc/Ori and NL4-3/Luc/KanR as obtained by inserting the low-copy bacterial origin of replication SC101 (Ori) or Kanamycin resistance gene (*Kan*R) in the pNL4-3/Luc backbone. **c,** Evaluation of transducing ability of lentiviral vectors as determined by luciferase activity of 293T cells transduced with VSV-G-pseudotyped NL4-3/Luc/Ori and NL4-3/Luc/KanR as compared to NL4-3/Luc, NL4-3/GFP and non-transduced 293T cells. **d,** Screening of 11 cell clones double-positive for NL4-3/Luc/Ori and NL4-3/Luc/KanR obtained by limiting dilution. For PCR, primers annealing to SC101 or KanR and pNL4-3 were used. **e,** PCR amplification of the LTR junctions from NL4-3/Luc/KanR/Ori double-positive cell clones a-e. **f,** Sequence analysis of a-e amplicons obtained in **e**. The red sequences indicate the T5 gRNA annealing site. Dashes indicate base deletions and green letters denote nucleotide mismatches compared to wild-type pNL4-3 LTR sequence. **g,** Schematic of inter-molecular concatemers bound in sense-sense orientation, as determined by nucleotide sequencing.

Five random colonies, named *A-E*, were picked and PCR checked for sense-sense inter-molecular concatemers as by Fig. 2e, and clones a-e, clearly distinguishable in the agarose gel of Fig. 3e, were sequenced at their LTR junctions (Fig. 3f). Sequence analysis confirmed that *a-d* clones were derived from excised NL4-3/Luc/Ori and NL4-3/Luc/KanR which were bound in sense-sense orientation as shown in Fig. 3g. Clones a and b contained large deletions, and clones c and d had a single nucleotide deletion and a single mutation compared to parental NL4-3 sequence, respectively. Parallel analyses performed with CRISPR/Cas9 and SC gRNA did not yield bacterial colonies (data not shown).

In all, these results demonstrate that NL4-3/Luc/KanR/Ori form spontaneously upon excision of NL4-3/Luc/Ori and NL4-3/Luc/KanR proviruses and that inter-molecular, sense-sense joining rebuilds full-length LTRs.

### HIV-1 concatemers show perceptible transcriptional activity but do not lead to infectious particles

To understand whether full-length LTRs in concatemers possess functional activity, 293T cells were transfected in parallel with NL4-3/Luc/KanR/Ori concatemers, pNL4-3/Luc, and pNL4-3/Luc/KanR. Three days after transfection, cells were divided in two, and one half was treated with CRISPR/Cas9 + T5 gRNA. Treated and untreated cells were cultured for five days and then analyzed for HIV *gag* mRNA expression, and for HIV proteins. Measurement of intracellular HIV *gag* mRNA was performed by real-time reverse transcriptase (RT)-PCR in three independent experiments. Compared to pNL4-3/Luc and pNL4-3/Luc/KanR, mRNA levels of NL4-3/Luc/KanR/Ori were much lower but detectable (Fig. 4a). As regards protein expression, pNL4-3/Luc/KanR and pNL4-3/Luc produced p24 levels that exceeded the highest limit of quantification of the ELISA used. These samples were therefore diluted 1:50 and retested. In all three experiments, NL4-3/Luc/KanR/Ori yielded results above the cut-off and comparable to pNL4-3/Luc/KanR and pNL4-3/Luc samples diluted 1:50. By contrast, pNL4-3/Luc/KanR cells treated with CRISPR/Cas9 + T5 gRNA produced p24 at levels below cut-off, further demonstrating deep HIV-1 impairment after DNA cleavage (Fig. 4b). Western blot analysis of cell lysates, performed with anti-HIV gag and IN, showed weak production of Gag and IN, confirming minimal transcriptional activity of NL4-3/Luc/KanR/Ori (Fig. 4c). Anti-Tat antibodies demonstrated Tat expression by pNL4-3/Luc and after giving Tat in *trans*, whereas no Tat can be detected after HIV concatemer transfection.

**Fig. 4.**
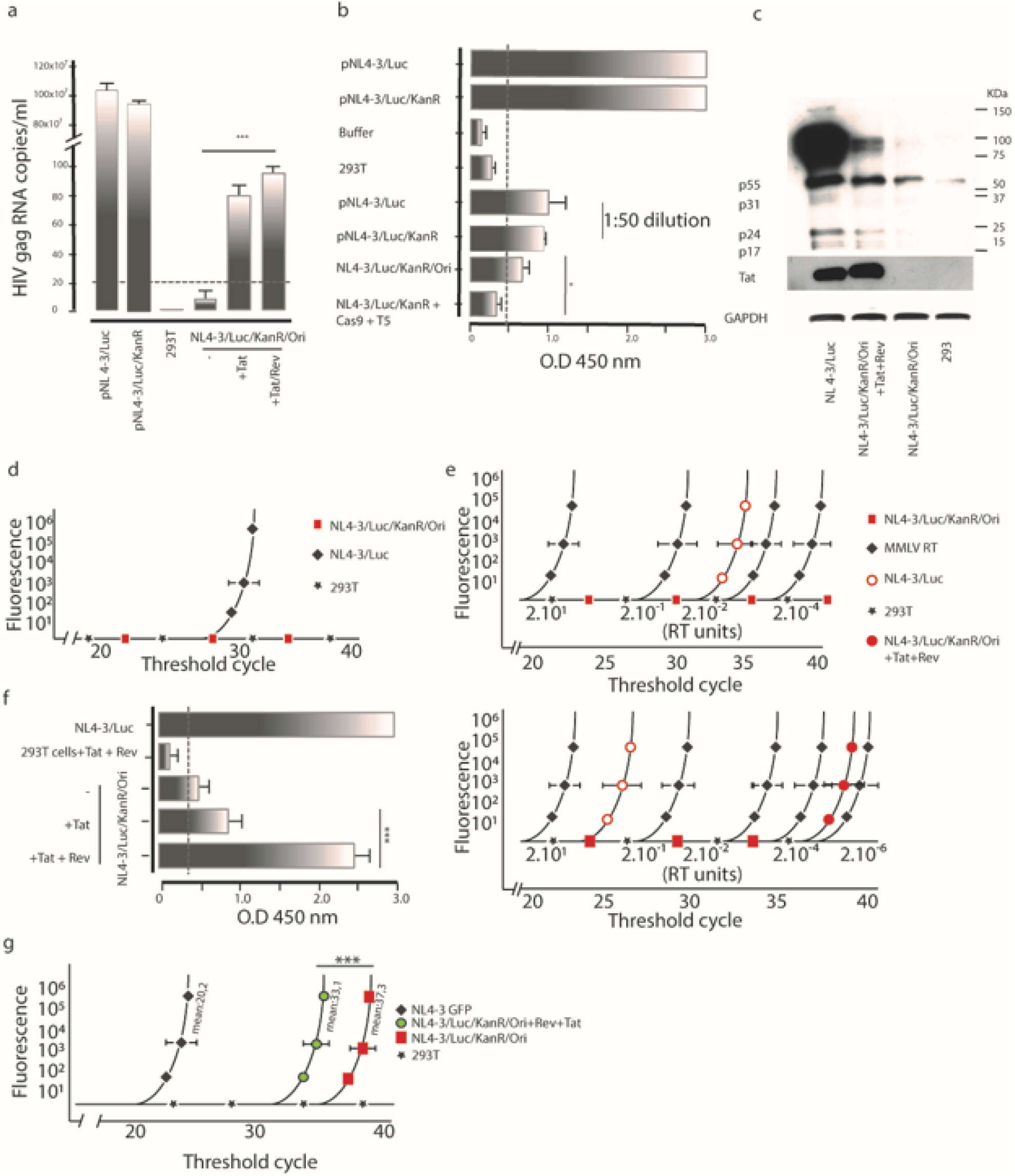
Without Tat and Rev, the concatemers are not transcribed. Analysis of HIV mRNA and protein production by NL4-3/Luc/KanR/Ori transfected alone or with Tat + Rev provided *in trans*. **a,** qRT-PCR quantitation of *gag* mRNA expression by cells transfected with pNL4-3/Luc or pNL4-3/Luc/KanR or pNL4-3/Luc/KanR/Ori alone or after transfection of Tat alone or Tat + Rev. Cells were lysed 3 days after transfection. Increment in *gag* mRNA expression following Tat or Tat + Rev provided in *trans* reached statistical significance (*p*=0.0013). **b,** Determination of p24 content by ELISA in lysates of cells transfected with pNL4-3/Luc or pNL4-3/Luc/KanR, as such or after 1:50 dilution, or with NL4-3/Luc/KanR/Ori before and after CRISPR/Cas9+T5 treatment. Production of p24 with pNL4-3/Luc and pNL4-3/Luc/KanR was comparable but significantly decreased following pNL4-3/Luc/KanR treatment with CRISPR/Cas9+T5. Isolated concatemers exhibited reduced but significant transcriptional activity. **c,** Western blot analysis performed on lysates of cells transfected with pNL4-3/Luc or NL4-3/Luc/KanR/Ori, either with Tat + Rev or alone, or not transfected 293T cells. Detection with antibodies anti-Tat, anti-p24, and anti-IN demonstrated low production of the three proteins by NL4-3/Luc/KanR/Ori that markedly increased after addition of Tat + Rev. **d,** TILDA, a method that detects multiply spliced RNA (msRNA), did not indicate production of msRNA by NL4-3/Luc/KanR/Ori-transfected cells. **e,** SG-PERT, which measures RT activity in supernatants as quantified by 10-fold dilutions of mammalian murine leukemia virus (MMLV) RT, was used to determine RT activity in supernatants of cells transfected with NL4-3/Luc/KanR/Ori or pNL4-3/Luc (upper panel) or NL4-3/Luc/KanR/Ori with or without Tat + Rev (lower panel). **f,** Determination of p24 content as in **b** and after transfection with Tat or Tat + Rev. **g,** Analysis of proviral DNA content of 293T cells cultivated in the presence of supernatants from cells transfected with pNL4-3/GFP, or NL4-3/Luc/KanR/Ori, alone or combined with Tat+Rev, or control 293T cells, and, for all, VSV-G provided *in trans*. This analysis, performed with HIV-Qual assay, demonstrated that addition of Tat + Rev, compared to NL4-3/Luc/KanR/Ori alone, yields increased HIV viral particle production. Data were collected from three independent experiments with four replicates each and analyzed by Student’s T test (*** = *p*<0.001).

To investigate whether expression of unspliced HIV-1 mRNA was accompanied and possibly driven by expression of multiply spliced HIV-1 mRNA (msRNA), we probed the cell lysates for *tat/rev* msRNA, a marker that has been shown to reflect the ability of a cell to produce virus^18,19^. To this aim, we used the *tat/rev* induced limiting dilution assay (TILDA), a method that discerns between latently and productively infected CD4^+^ T-lymphocytes^20^. As shown in Fig. 4d, and in contrast to pNL4-3/Luc, NL4-3/Luc/KanR/Ori did not express *tat/rev* msRNA. This result and the low p24 intracellular content suggested negligible viral replication. To further investigate this matter, we repeated the experiment by co-transfecting cells with NL4-3/Luc/KanR/Ori or pNL4-3 and VSV-G. At day 2 post-transfection, the supernatants were collected, assayed for the presence of RT activity, p24 and infectious particles. RT activity, tested using SYBR Green PCR-enhanced RT assay (SG-PERT)^21^, was negative for HIV concatemers alone (Fig. 4e, upper panel). As discussed below, provision of Tat + Rev in trans restored detection of RT activity in supernatants, even if at low levels (Fig. 4e, lower panel). In contrast, p24, assayed by ELISA, yielded an absorbance value slightly above the cut-off (Fig. 4f). The presence of infectious particles was examined by using these cell-free supernatants to transduce naïve 293T cells, that were then probed for the presence of NL4-3 provirus at day 3 post-transduction. This task was performed using Xpert HIV-1 Qual, a highly sensitive qualitative real-time PCR assay approved for *in vitro* diagnostics. In contrast to supernatants from cells treated with pNL4-3/Luc, which scored clearly positive (mean threshold cycle 20.2), the three replicas of NL4-3/Luc/KanR/Ori supernatants yielded weak amplification signals and reaching the threshold cycle at the 37^th^ cycle detection (Fig. 4g). Based on information provided by the manufacturer, experience accrued by routine use of the assay and reference papers^22, 23^, these values could not be scored as positive. These results confirm that inter-molecular concatemers are unable to produce infectious virions.

### Cells transfected with NL4-3/Luc/KanR/Ori release HIV-1 particles following addition of exogenous HIV-1 Tat and Rev

Prompted by previous reports showing that the HIV-1 genome persists for weeks as an episome in a latent form in lymphoid and myeloid cells and can be reactivated by superinfection^24–27^, we tested whether this may also occur with NL4-3/Luc/KanR/Ori. Since superinfection of NL4-3/Luc/KanR/Ori transfectants with another HIV virus would yield indistinguishable virions, 293T cells were transfected with the concatemer and, three days later, transfected again with HIV *tat* or *tat* + *rev* plasmids to mimic superinfection. Cells were monitored for intracellular HIV RNA (Fig 4a), p24 production (Fig. 4b and f) and virion release (Fig 4e) three days after the second transfection. As shown in Fig. 4a, provision of *tat* or *tat* + *rev* boosted *gag* RNA expression at levels significantly higher compared to those of cells transfected with NL4-3/Luc/KanR/Ori alone. Interestingly, in all three independent experiments, *gag* mRNA values were slightly higher in the presence of *tat* + *rev* compared to *tat* alone, suggesting that Rev, which exports unspliced HIV mRNAs from nucleus to cytoplasm, facilitates *gag* transcription. As expected, the presence of Rev had a more relevant boosting effect at a protein level: p24 content (Fig. 4f), which increased noticeably with Tat alone, reached levels similar to NL4-3/Luc in the presence of Tat and Rev. Fig 4c also shows that p55 increased proportionally in the presence of Tat + Rev (Fig. 4c).

To ascertain whether an increment in RNA expression and protein production corresponded to release of viral particles, NL4-3/Luc/Ori /KanR and *tat-* or *tat* + *rev-* transfected cells were further transfected with VSV-G. Supernatants from treated cells were tested for RT activity by SG-PERT two days later. As shown in Fig. 4e, lower panel, low but detectable RT activity in the supernatants was observed suggesting that Tat + Rev, provided in *trans*, triggered release of viral particles. To evaluate whether these were also infectious, supernatants (three replicas) were used to transduce 293T cells and these were examined for proviral DNA by Xpert HIV-1 Qual. Samples generated clearly positive amplification curves with a mean threshold cycle of 33.1 (Fig. 4g). This result indicates that 293T cells were transduced and, therefore, that the viral particles released following addition of Tat and Rev were infectious.

### HIV-1 IN activity facilitates persistence of the excised provirus in 293T cells

IN is thought to bind the 3’ ends of the linear cDNA of HIV and mediate integration of the proviral DNA in cell genome ^29,30^. Prompted by this, we hypothesized that the excised provirus mimics linear, non-integrated double stranded HIV DNA and, as such, can be concatemerized, thereby reconstituting two complete LTRs (Fig 2d). To address this hypothesis, we set up the experiment illustrated in Fig. 5a and opted to use GFP as a reporter gene. To mimic CRISPR/Cas9 cleavage, pNL-CMV-GFP, differing from pNL4.3-GFP because its heterologous protein expression is driven by the CMV promoter, was digested with *Pml*I, which has two blunt-ended cutting sites outside the HIV-1 genome. After digestion, the HIV-1 genome was extracted from agarose gel. As shown in Fig. 5a, the linearized HIV-1 genome was transfected alone or together with pCMV-IN, a plasmid encoding HIV-1 IN under the control of CMV promoter. Non-transfected cells and cells transfected with uncut pNL-CMV-GFP served as negative and positive controls, respectively. Three days post-transfection, cells were examined for GFP expression by flow cytometry and fluorescence microscopy. As shown in Fig. 5c and 5d, non-transfected 293T cells showed no fluorescence, whereas nearly 50% of the cells transfected with uncut pNL-CMV-GFP were fluorescent. As regards cells transfected with the linearized HIV genome, provision of HIV-1 IN *in trans* increased the percentage of fluorescent cells that, in three independent experiments, rose from about 3% to over 20% with an increment that was statistically significant (Fig. 5c and 5d). These experiments suggest that partial rescue of transcriptional capacity is facilitated by HIV-1 IN.

**Fig. 5.**
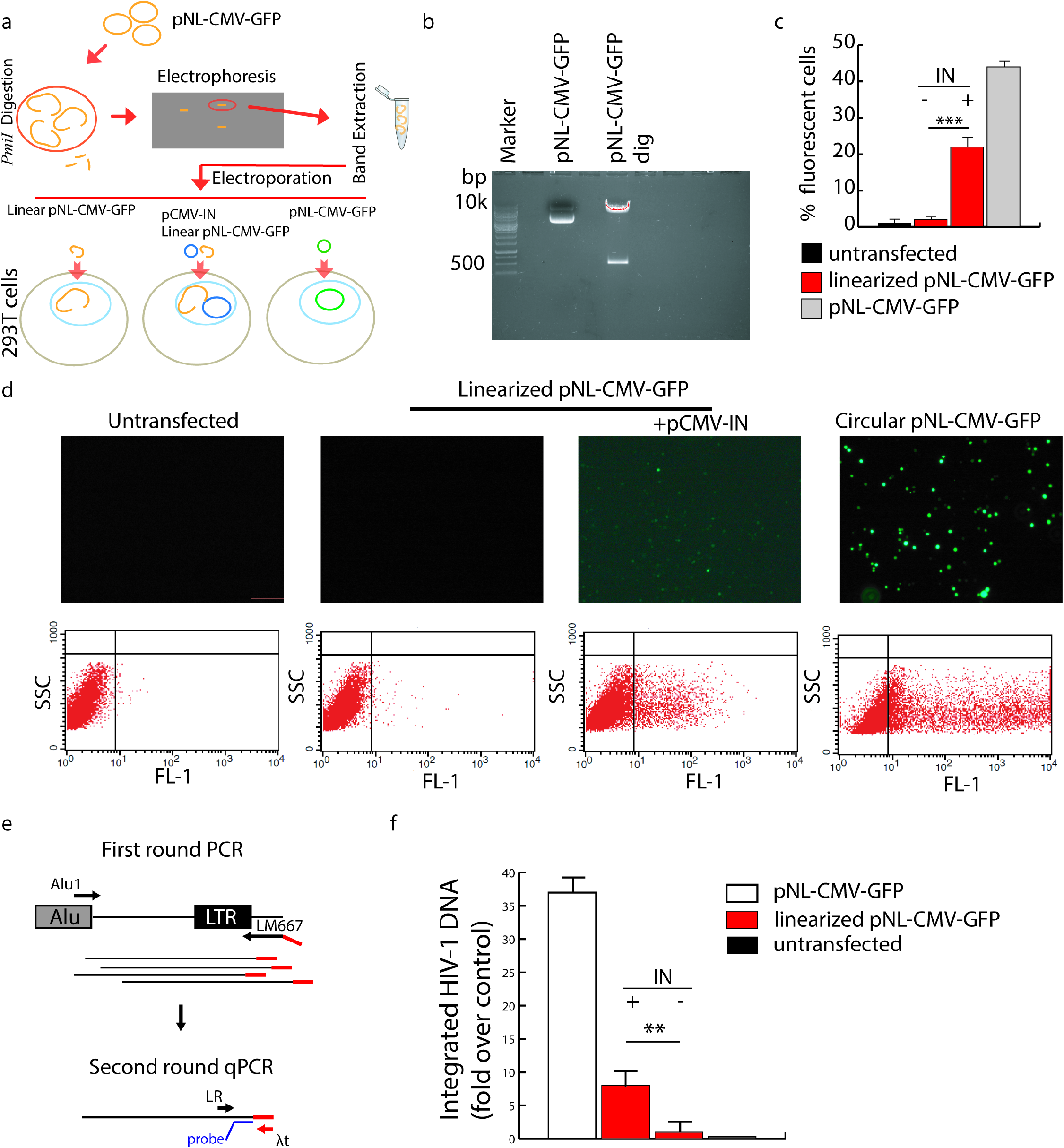
HIV IN contributes to circularization and integration of excised provirus. **a,** Schematic workflow of the experiment. The HIV provirus was obtained from pNL-CMV-GFP by digestion with *Pmi*I, gel purified and electroporated into 293T cells, either linearized, alone or in association to pCMV-IN, or as a closed plasmid. **b,** Agarose gel electrophoresis of pNL-CMV-GFP digested with *Pmi*I. The upper band denotes the linearized HIV provirus that was extracted, purified and used for transfection as described in **a**. **c,** Percentage of GFP+ 293T cells after transfection with pNL-CMV-GFP, linearized, with IN (+) or without IN (−), or closed, as determined by flow cytometry. Bars indicate SD as determined from three independent experiments. The percentage of fluorescent cells transfected with the linear provirus and pCMV-IN was significantly higher than for cells transfected with the linear provirus alone, as determined by Student’s T test (*p*<0.001). **d,** Fluorescence microscopy and flow cytometry analyses of the cells plotted in the histogram of Fig. 5c. Increment in cell fluorescence in the presence of HIV IN suggests that IN contributes to rescue transcriptional capacity of the linearized provirus. **e,** Schematic of Alu-PCR reaction. This reaction was used to understand if the provirus integrates into host cell DNA. The first round creates a tagged Alu-HIV-LTR fragment that is subsequently reamplified and detected by a specific probe. **f,** Histogram plot of the Alu-PCR reaction showing that the amount of provirus integrated back is significantly increased in the presence of HIV IN compared to the same without HIV IN. Statistical analysis was performed by one-way ANOVA and Bonferroni post-hoc test (p<0.001).

To understand if HIV-1 IN contributes to restoration of transcriptional activity by a novel HIV genome arising from the transcription of the excised and rearranged provirus, we took advantage of Alu repeated sequences interspersed in human genome to set up an HIV-1 Alu-PCR assay (Fig. 5e). Briefly, this assay employed a primer annealing to Alu and another one annealing to LTR sequences, and thus selectively amplified the HIV-1 genome integrated in Alu. The second primer contained the λt sequence tag, so that this PCR could be followed by a second one using a primer annealing to the sequence tag and another to LTR and a probe to quantitate the amplicons. HIV-1 Alu-PCR assay was run with DNA extracted from pNL-CMV-GFP - transfected and untransfected 293T cells (positive and negative controls, respectively), and 293T cells transfected with linearized pNL-CMV-GFP and further transfected or not with pCMV-IN. As shown by Fig. 5f, treatment with pCMV-IN increased the number of integrated pNL-CMV-GFP by about 7-fold compared to untreated cells. The increment was statistically significant. This experiment was performed in the presence of high amounts of linearized HIV-1 genome and HIV-1 IN, which could have facilitated interaction between molecules. Nonetheless, this experiment suggests that HIV-1 IN plays a role in novel integration events after gene-editing.

### The CRISPR/Cas9 excised HIV provirus persists and circularizes also in T-lymphocytes

With the aim to monitor the fate of the excised provirus in an *in vitro* model that closely mimics natural HIV infection, we repeated some experiments using Jurkat cells, a human T-lymphoid cell line, either actively producing or latently infected by HIV-1. The former was obtained by infecting cells with HIV NL4-3 strain. The latently infected cells were J-Lat cells, clone 9.2, harboring HIV-R7/E-/GFP, a full-length integrated HIV-1 genome expressing GFP. In such cells, HIV-R7/E-/GFP is present as a single copy per cell and persists in a latent phase from which it can be reawakened by treating cells with phorbol esters, tumor necrosis factor-α (TNF-α), or exogenous Tat^33^. For these features, J-Lat are widely used to study HIV latency and reactivation^34^.

Due to low sensitivity of T-cells to lipofection^35, 36^, J-Lat cells were CRISPR/Cas9 transfected using the ribonucleoprotein (RNP) nucleoporation system. For this purpose, we redesigned gRNAs to optimize efficiency of transfection. The first guide, named g1, targets the U5 region of LTR and the second, g2, binds the same R site targeted by T5 gRNA. As a negative control, we also designed a non-related gRNA (SC). To monitor formation of LTR circular molecules, we used droplet digital PCR (ddPCR) as described^37^. Primers and probe were designed in such a way to detect *nef*-LTR junctions, i.e. LTR circle molecules (depicted as green dots in Fig. 6a), as opposed to linear excised molecules that were not amplified (grey dots). Results were obtained from cellular DNA extracted 24 hours post-CRISPR/Cas9 treatment and normalized using primers and probe targeting housekeeping EIF2C1 gene (Fig. 6a).

**Fig. 6.**
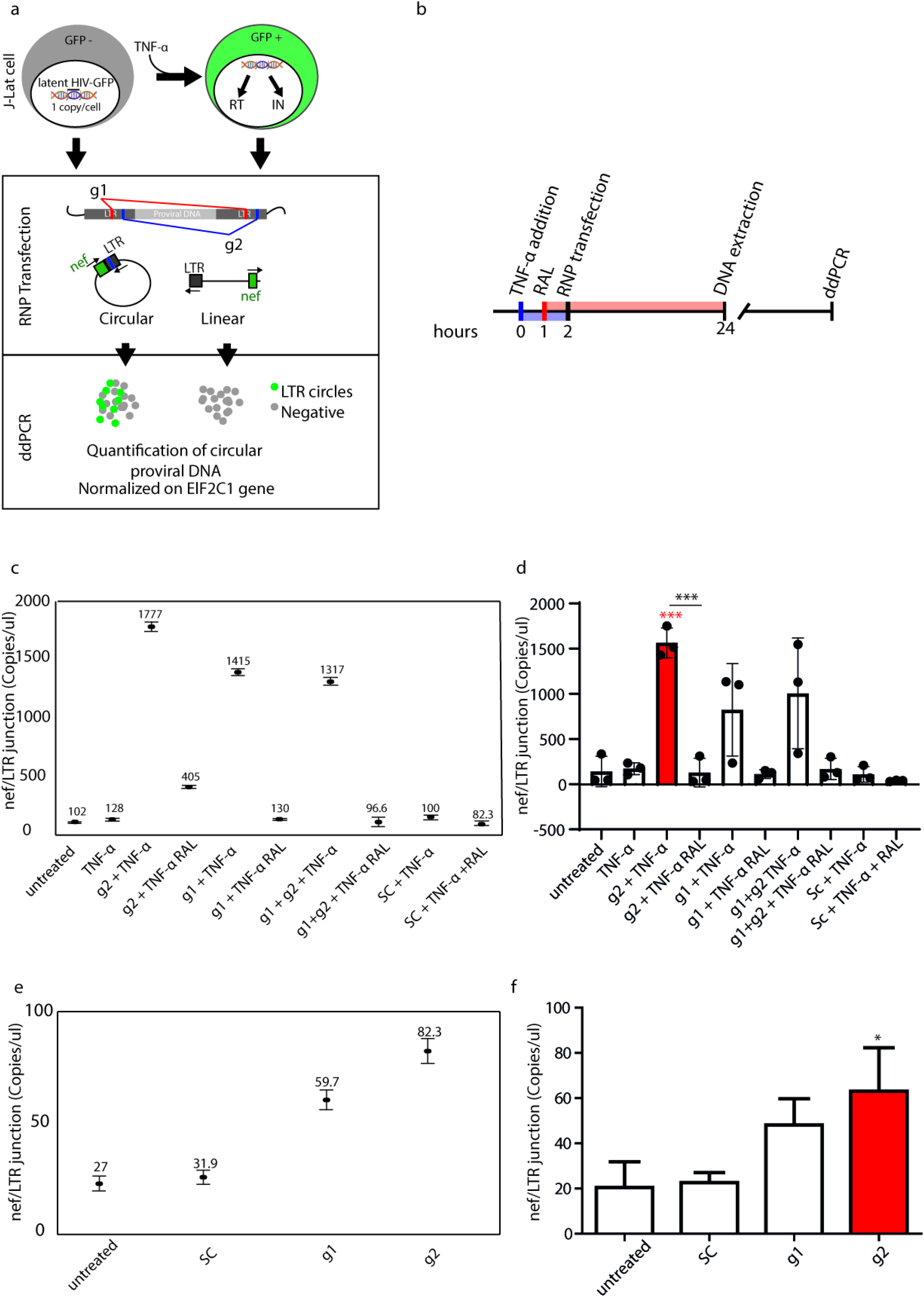
Circularization of the excised provirus is facilitated if cleaved out in the presence of HIV IN also in latently infected lymphocytes. **a,** Schematic illustration of J-Lat treatment, localization of g1 and g2 guides, and detection of LTR circles by ddPCR. J-Lat cells were exposed to TNF-α to activate HIV expression (and viral protein production) and then transfected with g1, g2, g1 + g2 or SC and incubated for 24 h as such, or with RAL. ddPCR was performed on genomic DNA digested with *Bse*JI using primers for HIV *nef* and LTR, that allow amplification of circles only. **b,** Timeline of drug addition, RNP transfection and total DNA extraction. J-Lat cells were treated with 10 ng/ml of TNF-α and 1 h later RAL, 10 μM, was added. RNP transfection was done after 2 h and, 12 h later, total DNA was extracted for ddPCR. **c,** A typical ddPCR experiment is shown, where *nef*-LTR junction concentration is determined. Results demonstrate that the number of LTR circle molecules significantly increased in cells treated with TNF-α. RAL treatment causes considerable decrease in LTR circle formation. Poisson error bars (confidence interval 95%) of ddPCR values are shown, as determined automatically by the software after every ddPCR run. **d,** Histogram plot of the average number of LTR circle molecules from 3 independent experiments (black circles), performed as described in **b**. Statistical analysis of ddPCR values, performed using single-way Anova (red asterisks) and Tukey’s multiple comparison test (black asterisks), shows that RAL pre-treatment decreased LTR circle molecule formation (*** *p* <0.001). **e,** The experiment described in **b** was performed without TNF-α or RAL treatment. A typical ddPCR experiment is shown as in **c**. Results demonstrate that LTR circle molecules are formed also in the absence of HIV activation. **f,** Histogram plot representation of 2 independent experiments as in **e**. Statistical analysis was performed using single-way Anova (* *p* <0.1).

Because HIV excision by CRISPR/Cas9 was conceived to cure latently infected cells and most studies were directed to target these cells ^3, 4, 7, 16^, we first investigated the fate of the excised provirus in J-Lat cells. As shown in the timeline of Fig. 6b, J-Lat cells were first treated with TNF-α and, after 1h, supplemented with 10 μM RAL, an HIV IN inhibitor. An additional hour later, we performed RNP transfection and total DNA was extracted 24 h after TNF-α addition. Fig. 6c shows the results of a typical ddPCR experiment: basal numbers of LTR circle molecules in J-Lat and J-Lat + TNF-α were below 100 copies. Similar numbers were found after treatment with CRISPR/Cas9 SC gRNA and TNF-α. In contrast, the number of HIV circular molecules increased 10-fold following TNF-α addition and CRISPR/Cas9 cleavage with g1 and g2 alone or in combination (Fig. 6c and d). These results show that not all gRNAs are equal. For reasons that were not addressed, the number of HIV circular molecules were consistently higher using g2, which targets the same LTR site as T5 (Fig. 6c). Interestingly, formation of LTR circle molecules upon CRISPR/Cas9 treatment occurred in the absence of TNF-α as well, although at a much lower concentration (Fig. 6e and f). These experiments indicate that circularization of the excised provirus also occurs in cells harboring one to very few numbers of integrated HIV genomes and circularization is enhanced by stimulating cells with TNF-α. The latter evidence implies other mechanisms, in addition to induction of HIV-1 transcription, that cause circularization of the excised genomes^32^ and take place in the absence of IN. These might be cellular proteins involved in NHEJ pathway.

### HIV IN helps reactivation of the HIV provirus in J-Lat cells

The observation that addition of TNF-*a* and editing promotes LTR circle molecule formation, together with our own findings (Fig 5) led us to hypothesize that the inhibition of IN might enhance CRISPR-Cas9 mediated eradication. To probe this idea, J-Lat cells, pretreated or not with 10 μM RAL for 24h, were then transfected with RNP with g1, g2, g1+g2 or the SC control guide labelled with Atto550. This dye allows tracking of RNP-transfected cells that appear fluorescently labeled in red. Five hours post-RNP treatment, HIV-1 transcription, and associated GFP expression, was induced by TNF-α, while keeping cells under RAL treatment. HIV reactivation was quantified in edited cells by enumerating GFP^+^Atto550^+^ cells by flow cytometry, performed 24 hours post-RNP treatment (Fig. 7a). Non-transfected, non-activated J-Lat cells had no red fluorescence and traces of GFP fluorescence (<1%, Fig. 7b, grey overlay histogram in every panel). TNF-α induction caused GFP^+^ cells to increase to 61.2% (Fig. 7b, dot plot, top panel). SC gRNA treatment did not affect HIV activation, as cells were nearly 70% GFP^+^ (Fig.7c, grey bar). Treatment with g1 or g2 reduced HIV activation by TNF-α to 40% (Fig. 7c, red bar) and 56% (Fig. 7c, blue bar) GFP+ cells, respectively. Again, we show that multiple targeting of LTRs has a stronger effect in reducing HIV activation compared to single targeting, since g1 + g2 proved to reduce GFP+ cells to 30% (Fig. 7c, green bar). A substantial percentage of cells were still GFP^+^ after RNP transfection. As shown in Fig. 7b and 7c, pretreatment with RAL reduced GFP expression over 10 times compared to RAL-untreated counterparts. Indeed, compared to 60% GFP^+^ cells observed in cell populations treated with SC gRNA and RAL, after treatment with g2, g1 and g1+g2, reduction was 91.7, 71.5 and 96%, respectively. These findings confirm the previous results on 293T cells, by demonstrating that IN plays a crucial role in relapse of viral transcription of gene editing in lymphocytes as well.

**Fig. 7.**
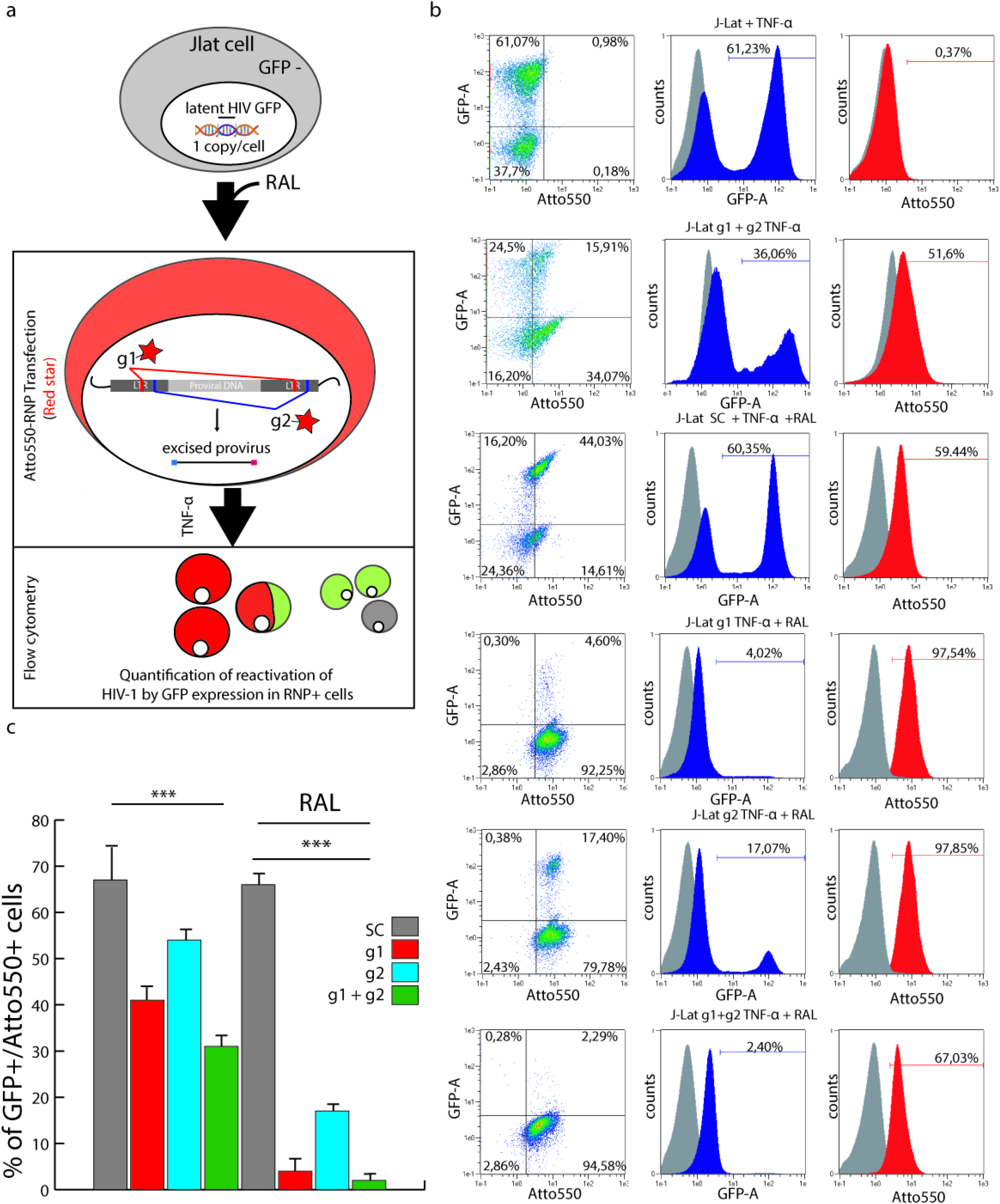
RAL pretreatment prevents reactivation of HIV after RNP transfection. **a,** Schematic workflow of the experiment: J-Lat cells, pretreated or not with 10 μM RAL, were transfected with Atto550-RNP containing g1, g2, g1+ g2, or the SC control guide. This allowed tracking of RNP-transfected cells that became fluorescently labelled in red. Five hours post-RNP treatment, HIV-1 transcription, and associated GFP expression, was induced by treating cells with TNF-α. HIV reactivation was quantified in RNP-treated cells by flow cytometry performed 24 hours post-RNP treatment. **b,** Flow cytometry analysis of J-Lat cells untransfected, transfected with g1, g2, g1 + g2, or SC, in the presence or not of RAL. All cells were activated with TNF-α 5 h after transfection. Ungated events are analyzed. The grey histogram in the overlays shows untreated, latent J-Lat cells. RNP transfection combined with RAL abolished HIV activation. **c,** Histogram plot of panels in Fig. 7b. The number of GFP^+^ cells is significantly lower upon RNP treatment with g1, g2 and g1 + g2 compared to SC gRNA. Combined treatment with g1 + g2 or with g2 alone is the most and the least effective, respectively, in reducing activated, GFP+ cells. Pretreatment with RAL together with RNP transfection dramatically reduces induction of HIV-1 transcription.

### IN and reverse transcriptase (RT) inhibition are essential for a more complete clearance of HIV-1 provirus

To test whether novel viral DNA could integrate in cellular genome in J-Lat cells after editing, we extracted J-Lat genomic DNA 24 h post-RNP treatment and probed it by Alu-PCR as described in Fig. 5e. Alu-LTR amplification was chosen so as to not amplify the single provirus integrated in J-Lat 9.2 cells, which has been mapped far from Alu sequences, i.e. within PP5 gene, chromosome 19, by two independent studies^39, 40^. Alu-PCR, therefore, did not amplify the original provirus but rather those that integrated back close to an Alu region. As shown in Fig. 8a, no integration in Alu was detected in latent J-Lat or J-Lat treated with any gRNA. Conversely, HIV activation by TNF-α increased the number of Alu-LTR elements, while integration was greatly reduced by RAL. This preliminary experiment shows that J-Lat cells are a suitable model to investigate integration by Alu-PCR.

**Fig. 8.**
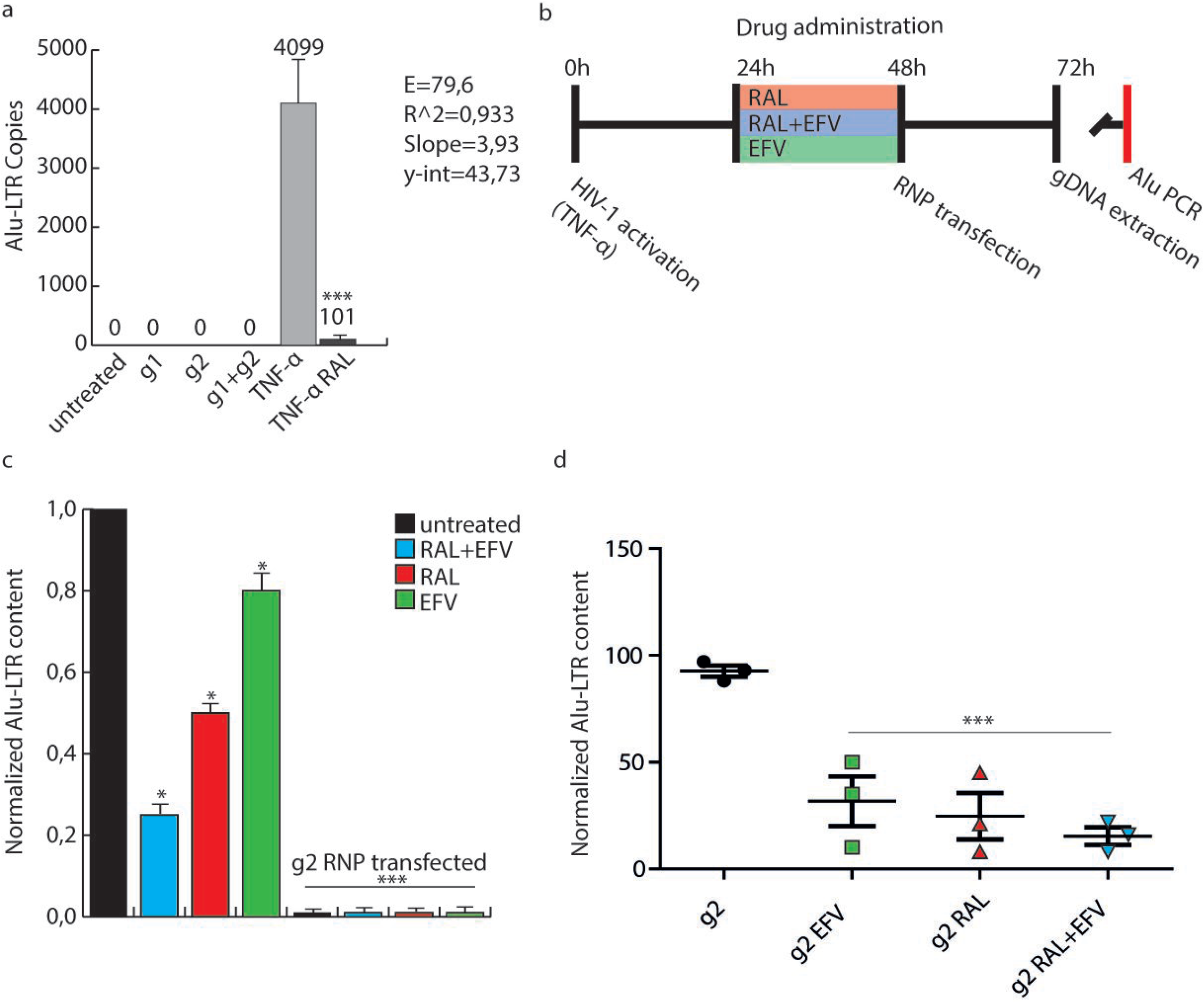
HIV-1 is reintegrated by IN and amplified by RT. **a,** Alu-PCR, performed as in Fig. 5, was carried out to detect the HIV-1 provirus integrated in proximity of Alu sequences of J-Lat cells. Alu-LTR copies are detected after activation with TNF-α and much less if treated with RAL. Latent cells treated with RNPs did not exhibit integration in Alu sites. **b,** Schematic workflow of the experiment shown in Fig. 8c and d. Cells were activated for 24 h with TNF-α, then treated with RAL or EFV or RAL + EFV for 24 h. RNP transfection with g2 was then carried out and the genomic DNA extracted 24 h later. Alu-LTR molecules were determined after normalization with β-globin DNA content. **c,** Histogram representation of Alu-PCR. RAL inhibits integration in Alu more than EFV alone, while combined treatment is more efficient than the single drugs alone. Guide g2 transfection greatly decreases Alu integration, whether alone or in the presence of RAL/EFV. **d,** Statistical analysis of Alu-LTR content of g2 RNP transfected cells alone. Single or combined treatments with RAL and/or EFV decreases efficiency of HIV reintegration. Data were obtained from three independent experiments and three biological replicates. Data were analyzed using One-way Anova (* *p*<0.1, *** *p*<0.001).

We therefore proceeded with the strategy exposed in Fig. 8b: briefly, cells activated for 24 h with TNF-α were treated with RAL or EFV, an HIV-1 RT inhibitor, for 24 h. RNP transfection with g2 was then carried out and, an additional 24h later, genomic DNA was extracted. Alu-LTR circles were determined after normalization with β-globin DNA content. Fig. 8c shows that: i) RAL inhibits integration in Alu more than EFV; ii) Combined treatment of RAL+EFV is more efficient than the single drugs alone in preventing integration; iii) most importantly, transfection with g2 alone dramatically reduces Alu integration in the presence of RAL/EFV. More detailed analysis of the differences among the g2-RNP transfected groups revealed that there is highly significant difference between the integration events occurring with or without associated drug administration, where RAL+EFV almost abolished reintegration in Alu. These data show that RT and IN play important roles in the number of HIV-1 proviral genomes integrated in Alu. Using both RAL and EFV after RNP transfection improved efficacy of HIV clearance.

### The CRISPR/Cas9 excised HIV-1 provirus circularizes and accumulates also in acutely wt HIV-1-infected lymphocytes

To assess whether cells acutely infected by wt HIV-1 accumulated circular HIV-1 genomes after editing, we repeated the study in Jurkat cells infected with a clinical isolate of HIV-1 at 0,05 MOI (Fig. 9a). Infected cells were treated with RAL and/or EFV for 7 days, then, CRISPR/Cas9 transfected using the same conditions described for J-Lat experiments. Two days after RNP transfection, we harvested culture supernatants and extracted total cellular DNA. Supernatants were examined for HIV genome content to determine reduction in viral replication following CRISPR/Cas9 treatment alone or supported by antiviral drugs. To evaluate whether CRISPR/Cas9 treatment increased the number of HIV-1 LTR circles in a model closely resembling natural HIV infection, cellular DNA was analyzed by ddPCR. As shown by Figures 9b and 9c, CRISPR/Cas9 transfection doubled the number of LTR circles in the absence of drugs (from 27.4 to 52.5 copies/μl). Similarly, the number of LTR circles was significantly higher upon CRISPR/Cas9 treatment in the presence of EFV, but not for RAL that was confirmed to decrease the number of circles. Interestingly, such increments, as well as absolute numbers of circle molecules, were much lower in the presence of both RAL + EFV, suggesting a synergistic effect (Fig. 9c and 9d). These findings recapitulate perfectly what observed with latently infected cells. Another striking similarity between acutely and latently infected cells is that RAL decreased the amount of LTR circles after HIV-1 ablation more than EFV, the RT inhibitor.

**Fig. 9.**
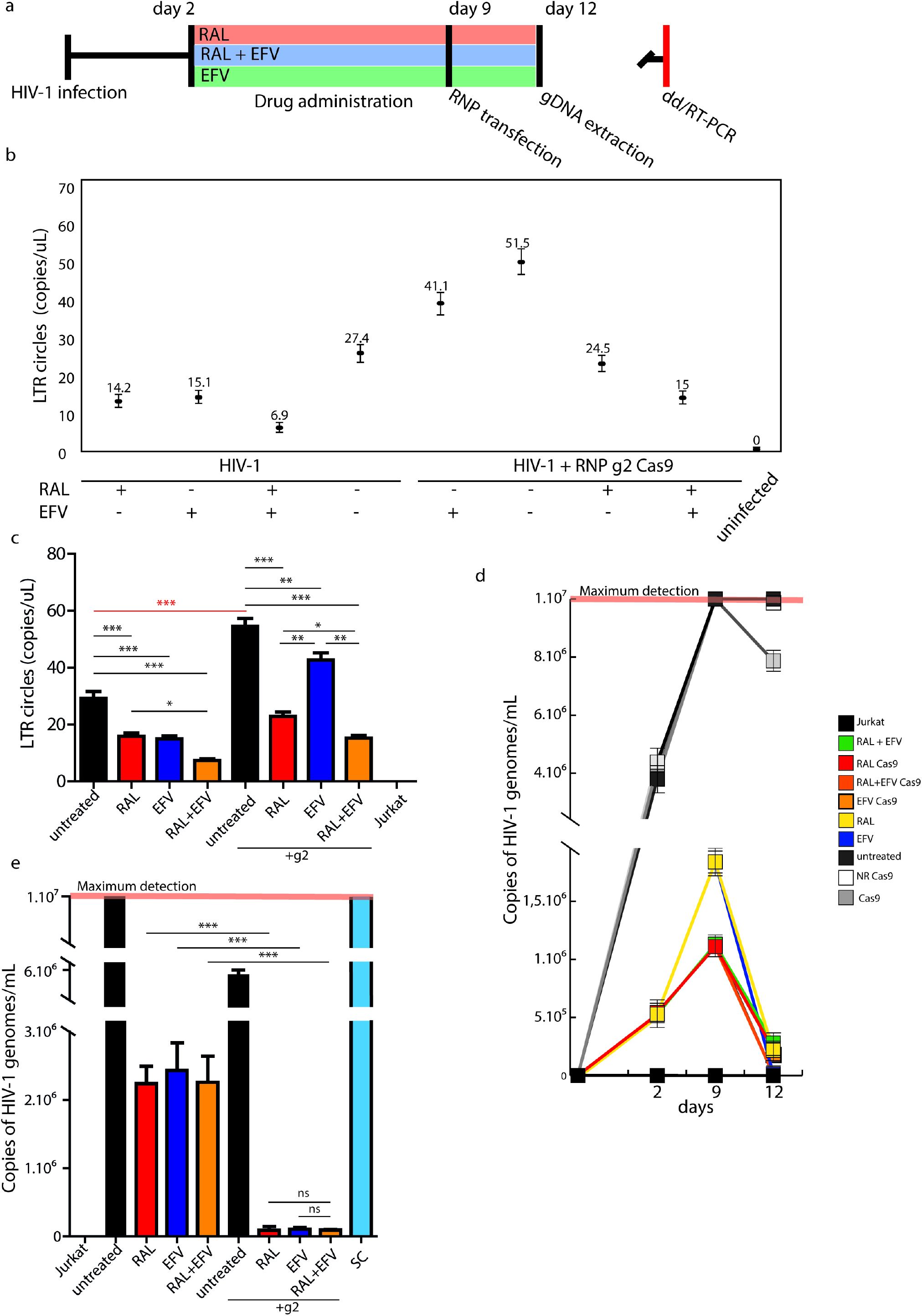
Cas9 proviral ablation increases the LTR circles of HIV-1. **a,** schematic illustration of the experimental procedure: 4 × 10^6^ Jurkat cells were infected with an HIV-1 clinical isolate. 24 h after the infection, cells were treated for 7 days with RAL 10 μM, EFV 100 nm and RAL 10 μM + EFV 100 nM. Then, cells were electroporated with Cas9 g2 RNP and processed for ddPCR. **b,** ddPCR shows an increase in LTR circles in Jurkat cells transfected with g2 RNP; RAL decreases the amount of LTR circles in both experimental groups (transfected/not transfected). Y axis = LTR circles concentration expressed as copies/μl. **c,** Statistical analysis performed on ddPCR reads. One-Way Anova with post-hoc Tukey test was performed, experiments are expressed as mean ± SD (* *p*<0.1, ** *p*<0.01, *** *p*<0.001). **d**, RT-PCR of viral RNA extracted from supernatants of cells described in **a** at 0, 2, 9 and 12 days from HIV infection. Y axis: copies of HIV-1 genomes/μl supernatant. **e**, Statistical analysis performed on RT-PCR reads at day 12. One-Way Anova with post-hoc Tukey test was performed, experiments are expressed as mean ± SD (* *p*<0.1, ** *p*<0.01, *** *p*<0.001).

Finally, we measured the change in viral load in time in supernatants from cells treated with antiviral drugs and CRISPR/Cas9. As shown in Fig. 9d and 9e, the combination of antiviral drugs and CRISPR/Cas9 transfection performed at d9 after infection reduced HIV viral load by 3 logs. This finding confirms the antiviral potential of CRISPR/Cas9 when associated with pre- and post-treatment with antiretroviral drugs.

## Discussion

CRISPR technology is becoming the leading gene editing tool, with increasingly expanding fields of application^11, 41^. These include HIV therapy, where CRISPR might allow excision of the integrated HIV genome, which can be excised from cellular DNA by taking advantage of CRISPR/Cas9 site-specific cleavage^42, 43^. The ends of cellular DNA are then joined by NHEJ^8^. This approach has proven to be effective and potent *in vitro*, whereas a number of limitations may be relevant when it is transposed *in vivo*. CRISPR/Cas9 may exhibit possible off-target activity and defiant gene rearrangement following DNA repair^41, 44^; moreover, diversity and mutations of HIV genome may constrain target selection^45–47^. Lastly, site of integration and transcriptional activity of the provirus may impact on susceptibility to CRISPR cleavage ^47–50^. *In vivo*, this scenario is further complicated by the lack of effective delivery vehicles^16^.

The main aim of this study was to monitor the fate of the HIV-1 provirus once it is excised, and the cellular repair mechanisms are triggered to heal the scars generated by CRISPR/Cas9. To this purpose, the experimental design was structured in two main parts. First, starting from previous observations showing that the HIV-1 provirus can be excised from 293T cell genome as in lymphoid T-cells^19, 20^, the fate of HIV-1 provirus was first studied in 293T transduced with a replication-deficient HIV-1-based vector. Second, findings were confirmed in a human T-lymphocyte cell line that contains one copy of integrated HIV-1 per cell and is a well-established model to study latency and reactivation^34, 35, 50^. Third, further confirmation was achieved in lymphoid cells infected with a clinical isolate of HIV-1.

One of the difficulties of CRISPR strategies is the control of the type of editing in cells and of the ensuing NHEJ process. Targeting the HIV-1 LTR alone, although allowing use of a single guide to cut out the provirus, has been repeatedly shown to rapidly lead to viral escape^5, 6, 7^. Thus, to effectively eliminate HIV infection *in vivo*, it would be advisable to convey multiple RNA guides into a single cell and ensure chopping of the provirus. Although rapid progress in CRISPR/Cas9 technology^41, 51^ and improvement of delivery strategies^52, 53^, will provide a way to circumvent these drawbacks, current technologies of *in vivo* delivery do not allow to control the number and amounts of RNA guides introduced into single cells. We found that, if cut with a single RNA guide, the excised HIV-1 provirus persists in cells for a protracted period. Depending on the number of copies per cell, it may persist in a linear form, with partially restored expression activity, but HIV-1 can also be found. The excised linear provirus differs from the linear HIV-1 cDNA produced by RT^12^ during viral replication *in vitro* and *in vivo* because it misses a portion of LTR at both ends. However, it may be converted into concatemeric HIV-1 cDNAs that either persist as episomes in infected cells or may be transcribed into a full viral genome that might follow the classic intasome-mediated integration pathway because, when they dimerize, they reconstitute the functional LTR^54^ at both ends. HIV episomes are believed to be the result of abortive integration processes^46, 64, 65^, deletion of sequences necessary for chromosome integration, or autointegration events occurring when an HIV cDNA integrates into another cDNA molecule^56^. They are found *in vitro*^57, 58, 59^ and in patients at advanced phases of the disease^13, 60–63^. Episomal HIV-1 genomes persist in lymphoid cells and macrophages suggesting that extrachromosomal HIV-1 and quiescent T lymphocytes are major reservoirs in infected individuals^63, 64, 67^. In keeping, episomal HIV-1, which also forms under antiretroviral regimens, has been postulated as a marker of ongoing *de novo* infection that may trigger viral rebound after treatment interruption^25, 68^.

Recent studies showed that episomes are neither rare nor inactive results of a dead-end process. Indeed, it has been shown that, despite the notion that integration is a prerequisite for protein expression, the HIV episome can be expressed. Expression of Tat and Nef or Tat alone by unintegrated HIV-1 activates resting T-cells and maintains persistent viral transcription in macrophages^14, 67^. Unintegrated HIV-1 is abundant in resting, non-proliferating CD4^+^ T cells and yields *de novo* virus production following cytokine exposure of resting cells^69^. Moreover, infection by HIV-1 mutants deleted of IN generated episomes producing Gag and Env *in vitro*, although they were uncapable to establish production of infectious HIV-1^7, 70^. Episomal DNA was found to express early gene msRNAs at a low level and, upon superinfection, it could also express late genes, indicating that it has the full potential for transcription^67, 70, 72^. Because of its potential for transcription, the unintegrated DNA may influence viral RNA decay consequent to therapy, and even recombine with a second incoming virus, thus contributing to the generation of viral diversity^26, 27, 68^.

The results of our study support the observation that CRISPR/Cas9 gene editing is an extremely powerful technique that allows HIV excision. However, our work demonstrates that CRISPR/Cas9 excised provirus persists in cells and can re-enter the replication flow. In transduced 293T cells, we found that two or more linear fragments could bind together in sense-sense orientation and express unspliced mRNA at low levels. Importantly, as described elsewhere for non-integrated HIV DNA^24, 67, 70, 72^, this viral DNA responded to exogenous Tat and Rev. Tat alone remarkably increased the level of expression of unspliced mRNA but, when tested at a protein level, Rev made the real difference, as it significantly augmented the amount of p24 protein. Further, infectious VSV-G pseudotyped virions could be detected in these supernatants. This indicates that provision of exogenous Tat and Rev also promotes release of viral particles from HIV episomes. It might be argued that transfection of Tat and Rev, chosen to facilitate measurement of concatemer products, led to much higher protein levels compared to superinfection, and that this is a rare event *in vivo*. How rare it really is remains still poorly defined. Besides superinfection by a second strain, which is indeed rare, Geldelblom and colleagues showed that the same HIV strain can reinfect the cell, a process known as co-infection^24^.

Interestingly, the fate of excised provirus was influenced by the number of provirus copies per cell. Indeed, RT activity increased the amount of HIV-1 provirus transcribed from RNA in activated J-Lat cells, possibly raising the chances of dimer/concatemer formation and reintegration (Fig. 8). Concatemer formation reconstituting 2 functional LTRs was detected after CRISPR in 293T cells. This might be due to the fact that multiple copies of provirus are present, a condition not so rare and that has been observed *in vivo^82^*. For example, when HIV-1 infection spreads through virological synapses, i.e. adhesive structures between infected and uninfected cells, multiple copies of HIV-1 are transmitted to the engaged uninfected cells^81, 83–86^. This phenomenon occurs in experimental models and in humans^87–89^, and has been linked to reduced sensitivity to antiretrovirals^82, 85, 90^. To understand what happens in a less artificial system, we repeated the experiments in J-Lat cells, an HIV-1 infected T-cell line harboring one HIV copy per cell, which persists in a latent state, and, finally, in Jurkat cells infected with a clinical isolate of HIV-1. As in 293T cells, excision through HIV-LTRs determined a sharp and statistically significant increment in episome numbers. In J-Lat cells we found that, in the absence of inhibition of RT and IN, the integration events are more abundant.

IN seems to exert a critical role in rescuing proviral DNA, a fact that is not surprising; IN takes part in various steps of HIV replication^29, 30^. Integration itself is a complex, multistep process catalyzed by IN that inserts the viral DNA ends into the cellular DNA strands. Integration creates a 2-nucleotide gap in the viral genome^30^ that is then repaired by cellular DNA repair machinery after integration^73^. Screening of knockout libraries showed that NHEJ plays a chief role in this process, and depletion of some cellular proteins involved in this pathway decreases provirus integration and viral infectivity^74–77^. It has also been demonstrated that some NHEJ proteins bind IN and this complex recruits the catalytic subunit of DNA-dependent protein kinase which triggers a cascade of phosphorylation and fills the nucleotide gap^78–80^. From our results, it can be speculated that activation of NHEJ by CRISPR/Cas9 facilitates circularization of excised proviral DNA. Although this mechanism was not thoroughly investigated, in both cellular models studied, circularization and reactivation could be enhanced or prevented by adding exogenous IN or inhibiting it, respectively. No matter what cells were used, IN played a crucial role in circularization and HIV persistence, as demonstrated by the reduction in LTR circle molecule formation and integration in cells treated with the IN inhibitor RAL. Interestingly, in a recent paper, Dash and colleagues treated HIV-1 infected humanized mice with a long-acting slow-effective release antiviral therapy for 4 weeks and three weeks later with CRISPR/Cas9 to excise the provirus. The drug regimen included an IN inhibitor. Five out of 7 mice showed rebound of viremia at levels comparable to animals that received no CRISPR/Cas9 treatment. The Authors did not investigate why this occurred but speculated that, in the 5 animals, the virus rearranged to reinstate competence for replication^4^. In the light of our results, it would be informative to perform CRISPR/Cas9 treatment in the context of antiretroviral treatment that includes RAL, to shed light on the role of IN and RT in shaping the fate of the excised provirus *in vivo*. We did not observe any difference between LTR circles in J-Lat cells before or after TNF-α administration, showing that activation alone cannot increase the number of circles detected (Fig. 6d).

Interestingly, circles are indeed present also in Jurkat cells, no matter whether the virus is productively replicating or dormant. These data show that circles do form without activation, even if at an obviously much lower number. These data suggest that cutting DNA, thereby activating NHEJ, brings to the formation of LTR circles. Activation determines amplification of this phenomenon by the combined activity of RT and IN (Fig. 6 and 8).

In conclusion, we provide evidence that if the HIV-1 genome is excised as a single fragment, it persists and reorganizes in concatemers giving the virus a second chance to express itself and rebuild an infectious form. Most concatemers and episomes are likely to be lost during subsequent cell mitosis and have limited persistence in dividing cells but are nonetheless a warning note. This work stresses the importance of CRISPR/Cas9 strategy in the cure of HIV, and should be a stimulus to 1) implement the efficacy of delivery systems and CRISPR/Cas9 strategies *in vivo* to achieve cleavage of HIV genome in multiple sites and in all cells, no matter where the provirus is integrated and how many copies are present within the cell; 2) prolong antiretroviral treatment to allow excised provirus to be eliminated.

## Materials and Methods

### Cell cultures and plasmids

Human 293T cell line was purchased from American Type Culture Collection (ATCC) and cultured in Dulbecco’s Modified Eagle’s Medium (DMEM) supplemented with 10% fetal bovine serum (FBS), 2 mM L-glutamine and antibiotics (penicillin, streptomycin) at 37°C and 5% CO_2_. The Jurkat T-cell line was purchased from ATCC and grown in RPMI 1640 supplemented with 10% FBS, 2 mM L-glutamine and antibiotics at 37°C and 5% CO_2_ and infected with a clinical isolate of HIV-1 at a multiplicity of infection (MOI) of 0.05. The human T-lymphoid J-Lat cell line was obtained through the NIH AIDS Reagent Program, Division of AIDS, NIAID. J-Lat cells were produced by transducing Jurkat cells with HIV-R7/E-/GFP at low multiplicity of infection (MOI) in such a way to generate clones containing one copy of integrated HIV per cell. Cells were cultured in RPMI 1640 supplemented with 10% FBS, 2 mM L-glutamine and antibiotics at 37°C and 5% CO_2_. J-Lat HIV-R7/E-/GFP, which is full length HIV-1 genome with a non-functional Env due to a frameshift, and GFP in place of the Nef gene, generates incomplete virions. HIV-R7/E-/GFP is activated for transcription and expresses GFP by treating J-Lat with TPA, TNF-α, or exogenous Tat^34^. Activation was performed by supplementing J-Lat cells culture medium with 10 nM TNF-α.

pNL4-3/Luc (https://www.aidsreagent.org/pdfs/ds3418_010.pdf Cat n: 3418) and pNL4-3/GFP (https://www.aidsreagent.org/11100_003.pdf, Cat n:11100) were obtained through the AIDS reagent program. The first is an HIV-1 NL4-3 luciferase reporter vector that contains defective Nef, Env and Vpr; it is competent for a single round of replication. It can only produce infectious virus after cotransfection with env expression vector. The second is also derived from pNL4-3 but carries enhanced green fluorescent protein (EGFP) in the *env* open reading frame. This vector expresses an endoplasmic reticulum (ER)-retained truncated Env-EGFP fusion protein. For linearization experiments (Fig. 5) we used plasmid pNL-EGFP/CMV/WPREdU3 (pNL-CMV-GFP) (Addgene, MA, USA). pNL4-3/Luc/Ori and pNL4-3/Luc/Kan were produced by molecular cloning into pNL4-3/Luc. Both SC101, the low copy bacterial origin of replication, and *KanR* were extracted from pSF_CMV-SC101 (#OG13, Oxford Genetics, Oxford, UK) following digestion with *Swa*I (SC101) and *Pme*I (*KanR*). Fragments were cloned in *BseJ*I site of pNL4-3 *env*.

### Digestion of host and linear DNA

Host and proviral DNA were extracted from cells using a standard phenol/chloroform method. Briefly, we added one volume of phenol:chloroform:isoamyl alcohol (25:24:1) per sample and vortexed for 20 seconds, then centrifuged for 5 min at 16,000 × *g*. The aqueous phase containing total DNA was purified by ethanol precipitation. Three μg of purified DNA were treated with 2 μl of Plasmid-Safe ATP-dependent exonuclease (Epicentre, Madison, USA) at 37°C for 30 minutes and then heat-inactivated by a 30-min incubation at 70°C.

### RCA

Following Plasmid-Safe ATP-dependent exonuclease treatment, circular DNA molecules were amplified with random examers and TempliPhi DNA polymerase (Merck KGaA, Darmstadt, Germany) following manufacturer’s instruction. Briefly, the reaction mixture containing 10 ng DNA was incubated at 30°C for 6 h. The reaction was then blocked by heat inactivation at 65°C for 5 min.

### CRISPR/Cas9 design and transfection

T5 (TTAGACCAGATCTGAGCCT), the CRISPR/Cas9 gRNA targeting the LTR R region and used in 293T cells, was cloned into pSpCas9-2A-Puro (Cat. No. 62988, Addgene, MA, USA) or in pU6-Cas9-T2A-mCherry (Cat. No. 64324, Addgene) following a standard protocol^92^. Transfection of DNA plasmids into 293T cells was performed with standard calcium phosphate method. J-Lat cells were transfected with CRISPR/Cas9 RNP and tracr-RNA Atto550 (IDT, Coralville, Iowa) by electroporation (NEON Electroporation System, Thermo Fisher, Massachusetts, USA) using the following parameters: 1400 V, width 10 ms, 3 pulses. Guides 1 and 2 (g1 and g2) were designed using IDT algorithm and as follows: g1 - /AlTR1/rUrGrArCrArUrCrGrArGrCrUrUrUrCrUrArCrArArGrUrUrUrUrArGrArGrCrUrArUrGrCrU/AlTR2/; g2 –/AlTR1/ rArCrUrCrArArGrGrCrArArGrCrUrUrUrArUrUrGrGrUrUrUrUrArGrArGrCrUrArUrGrCrU/AlT R2/; both gRNAs target LTR, g2 the same R site as T5, g1 targets U5.

### 293T cell transduction

. Transduction of 293T cells was performed with VSV-G-pseudotyped particles. These were produced by transfecting 293T cells with pNL4-3 or its derivatives described above and VSV-G plasmid. Generated particles were harvested at day 2 or 3 post-transfection and titrated as described elsewhere^93^. Transduction was performed using MOI 5 to transduce 2 × 10^5^ 293T cells. Cloning of transduced 293T cells was performed in 96-well plates at the indicated days post-transduction.

### ddPCR

For ddPCR analysis, total DNA from 5 × 10^5^ J-Lat cells was purified using Qiagen blood mini kit (Qiagen, Hilden, Germany) and quantitated spectrophotometrically. Aliquots of 2.5 μg were then *BseJ*I digested (Thermo Fisher, MA, USA) to fragment the genomic DNA without cutting the LTR junctions. Restriction products were column-purified using PCR Clean Up kit (Qiagen) and 2.5 ng of purified DNA were amplified by ddPCR using the following primers: LTR circles: Fwd - AACTAGGGAACCCACTGCTTAAG; Rev TCCACAGATCAAGGATATCTTGT;

Probe (FAM) – ACACTACTTGAAGCACTCAAGGC. Reaction was performed as follows: 10 min at 95°C denaturation, 40 cycles of 95°C for 30 s, and 60°C for 60 s, followed by 98°C for 10 min. After completion of PCR cycling, reactions were placed in a QX200 instrument (Bio-Rad, Milan, Italy) and droplets analyzed according to manufacturer’s instructions. Normalization was performed targeting the EIF2C1 gene, using a premade probe solution (Bio-Rad) following manufactures’ instruction.

### Concatemer isolation, characterization and sequencing

Selection of clones double positive for pNL4-3/Luc/Ori and pNL4-3/Luc/Kan was performed by PCR amplification using the followin g primers: Env Fwd – GACACAATCACACTCCCA; Kan Rev - AATAGCCTCTCCACCCAA; Ori Rev – TGTGGTGCTATCTGACTT. Selected clones were then transfected with T5 gRNApspCas9 plasmid (Cat. No. 459 Addgene) and cultivated in the presence of Puromycin (3 μg/ml) to select for the transfected cells. Following DNA extraction and digestion with ATP-dependent exonuclease, 50-200 ng of total DNA, quantitated before exonuclease digestion, were used to transform 10^8^ ultracompetent Stbl2 *E. coli* cells (Invitrogen, Carlsbad, CA, USA). Transformants were seeded in Luria Bertani (LB) agar plates supplemented with Kanamycin (50 μg/ml). Bacterial colonies were picked and screened to eliminate spurious clones containing residual pNL4-3/Luc/Kan, which also harbors the Ampicillin resistance gene in the plasmid vector. To this aim, colonies were split and seeded in two LB broth cultures containing Kanamycin or Ampicillin. Clones growing only in Kanamycin medium were expanded and concatemers extracted and purified using MAXI prep (Qiagen). Recovered DNA was tested by PCR to confirm the LTR junction, using the following primers: Fwd - AACTAGGGAACCCACTGCTTAG; Rev - GACAAGATATCCTTGATCTGTGGA. Positive clones were sequenced in LTR junctions by cycle sequencing.

### Biological activity of excised proviruses and concatemers

Analyses were focused on the detection and quantitation of HIV mRNAs and proteins and performed on cell lysates obtained by transfecting 1 × 10^5^ 293T cells with 0.5-1.0 μg of pNL4-3 derivatives. The same analyses and infectivity as the released virions were assessed with 1 × 10^5^ 293T transfected with 0.5-1.0 μg pNL4-3 derivatives and 0.1 μg VSV-G plasmid. Total cellular RNA was extracted using Maxwell 16 LEV simply RNA extractor (Promega, Madison, USA). Total proteins were extracted using RIPA buffer (0.22% Beta glycerophosphate, 10% Tergitol-NP40, 0.18% Sodium orthovanadate, 5% Sodium deoxycholate, 0.38% EGTA, 1% SDS, 6.1% Tris, 0.29% EDTA, 8.8% Sodium chloride, 1.12% Sodium pyrophosphate decahydrate).

### Measurement of HIV RNA and HIV DNA

Intracellular and supernatant HIV RNA were quantitated using COBAS AmpliPrep and COBAS-6800 (Roche, Milan, Italy), respectively. Both platforms and tests are routinely used at the Virology Unit, Pisa University Hospital, are certified for *in vitro* diagnostics and detect up to 20 HIV RNA copies/ml. HIV msRNA was detected and quantitated using TILDA^20^. Briefly, total RNA extracted as above was reverse transcribed at 50°C for 15 min, denatured at 95°C for 2 min and amplified for 24 cycles (95°C 15 seconds, 60°C, 4 min) on T100 PCR instrument (Biorad, Hercules, CA, USA). At the end of this process, samples were diluted to 50 μl with Tris-EDTA buffer and 1 μl of sample was used as template for a second *tat*/*rev* real-time PCR reaction. Primer sequences and details to calculate mean and standard deviation of ΔCT (threshold cycle) are provided elsewhere^20^. Proviral DNA in 293T cells was assayed with Xpert HIV-1 Qual, manufactured by Cepheid (Milan, Italy) and certified for *in vitro* diagnostics. This assay has a sensitivity of 278 copies/ml in whole blood^22, 23^. Before Xpert HIV-1 Qual analysis, genomic DNA extracted from 293T cells was RNase treated to eliminate contaminating cellular RNA.

### HIV-1 integration assay

(Alu-PCR). HIV-1 integration was examined in 293T cells and J-Lat cells by isolating genomic DNA from 1 × 10^6^ cells using the DNeasy Tissue Kit (Qiagen). Alu-LTR sequences were amplified during first round PCR from 100 ng of total genomic DNA using the following primers: LM667 5’-ATGCCACGTAAGCGAAACTCTGGCTAACTAGGGAACCCACTG; Alu1 5’-TCCCAGCTACTGGGGAGGCTGAGG. Amplification was performed for 16 cycles performed as follows: denaturation for 3 min at 95°C, 16 cycles of 95°C for 30 s, and 60°C for 60 s, followed by 72°C for 60 s. First round PCR is followed by a second round real-time PCR using a 1:500 dilution of first PCR mixture together with the Alu-specific primer λT (5’-ATGCCACGTAAGCGAAACT),U5 LTR primer LR (5’-TCCACACTGACTAAAAGGGTCTGA) and probe ZXF-P81 (5’-FAM-TGTGACTCTGGTAACTAGAGATCCCTCAGACCC-TAMRA). Real-time PCR amplifications were performed on a CFX96 machine (Bio-Rad). Results were normalized using the single-copy Lamin B2 gene that was quantified by real-time PCR (Fwd 5’-CCCCAGGGAGTAGGTTGTGA; Rev 5’-5′-TGTTATTTGAGAAAAGCCCAAAGAC).

### Western blot and HIV protein detection

Total proteins were extracted by direct lysis of samples using RIPA buffer. Extracted proteins were titrated using the Bradford assay and then analyzed by ELISA or Western blot. Capsid p24 was measured in cell extracts and supernatants using SimpleStep ELISA (ABCAM, Cambridge, UK) and ADVIA Centaur HIV Ag/Ab Combo ELISA (Siemens Healthcare Diagnostics, NY, USA), respectively. RT activity in supernatants was determined by SG-PERT and as described by Vermeire and colleagues^94^. Western blot analysis was performed using 20 μg RIPA-extracted proteins and using a mixture of antibodies anti-Tat, anti-Gag, and anti-IN (ab42359, ab63917 and ab66645, AbCam, Cambridge, UK) or anti-GFP (ab183734, Abcam), or anti-actin (ab179467, Abcam).

### Flow cytometry

Fluorescent cells were measured using Attune NTX (Thermoscientific, USA) or FACScan (Becton-Dickinson, Florence, Italy). Cells were analyzed 2-3 days post-transfection or as indicated, following detachment from well plates by Trypsin treatment and pelleting by centrifugation at 300 × *g* for 5 min. Data were analyzed using FSC Express 4 software (DeNovo Software, Glendale, CA).

### Statistical analysis

GraphPad Prism software V5.03 (GraphPad Software, Inc., USA) was used for statistical analysis. Data were analyzed using the Student’s *t*-test. Differences between groups were considered statistically significant at values of *p* < 0.05. All results, including flow cytometry and ddPCR and Alu-PCR data, were obtained from at least three independent experiments and expressed as mean ± standard error of the mean (SEM). ddPCR and Alu-PCR results were analyzed using one-way ANOVA, **p < 0.01, n.s. not significant. Flow cytometry assay results were analyzed using the Student’s *t*-test, *** p<0.001, ** p<0.01, * p< 0.1.

## Acknowledgements

This work was supported by “*SENSOR, nuovi sensori Real-Time per la determinazione di contaminazioni chimiche e microbiologiche in matrici ambientali e biomedicali”*, Progetto co-finanziato dal POR FESR Toscana 2014-2020; “*Addressing viral neuropathogenesis: Unraveling the molecular and cellular pathways of viral replication and host cell response and paving the way for the development novel host-targeted, broad spectrum, antiviral agents*”, bando PRIN: Progetti di ricerca di rilevante interesse nazionale, Bando 2017, Prot. 2017KM79NN; and “*I-GENE, In-vivo Gene Editing by Nanotransducers*”, European call identifier H2020-FETOPEN-2018-2020, Proposal ID 862714.

## Author contributions

ML, EM, OT, GA, and MP conceived and designed the experiments; ML, PQ, GM, SC, SP performed and analyzed the experiments; FM, MS, GF, OT, JLH supervised analyses, and provided reagents and resources, MP supervised experiments and provided funding; ML, GF and MP wrote the manuscript with input from all other authors.

